# Condition-dependent, amorphous protein agglomerates control cytoplasmic rheology

**DOI:** 10.1101/2025.06.17.660151

**Authors:** José Losa, François Simon, Dmitrii Linnik, Saniye Gül Kaya, Marc C. A. Stuart, Artem Stetsenko, Rinse de Boer, Franz Y. Ho, Danny Incarnato, Jan Stevens, Jan van Eck, Marco W. Fraaije, Lucien E. Weiss, Sven van Teeffelen, Sanne Abeln, Albert Guskov, Siewert-Jan Marrink, Bert Poolman, Matthias Heinemann

## Abstract

Molecular crowding in the bacterial cytoplasm restricts the diffusion of large molecules, impacting cellular processes. However, how nutrient availability influences cytoplasmic rheology is not well understood. With single-particle tracking in *Escherichia coli*, we observed a threefold variation in the diffusion of a 40-nm particle across exponential growth conditions. Previously suggested determinants of rheology did not account for this variation; instead, we found a strong anticorrelation between the diffusion coefficient and the abundance of amino acid metabolism proteins, persisting upon genetic perturbations and showing that lower diffusion is associated with increased viscoelasticity. Photoactivated light microscopy revealed that some amino acid metabolism proteins form clusters. Electron microscopy showed that these proteins could form amorphous agglomerates at physiological concentrations in vitro, likely driven by their low intrinsic disorder, high compactness and hydropathy score. These findings show that protein agglomerates regulate cytoplasmic rheology in a condition-dependent manner, suggesting an underappreciated level of cytoplasmic organization.

**Highlights:**

- Diffusion of 40-nm particles varies threefold across growth conditions in *E. coli*
- Cytoplasmic diffusion inversely correlates with COG-E protein abundance COG-E proteins form agglomerates that increase cytoplasmic viscoelasticity
- Protein compactness and hydrophobicity predict condition-dependent crowding effects

## Introduction

The cytoplasm is a densely packed environment where, in bacteria, macromolecules such as proteins and RNA reach concentrations of approximately 300 g/L and 100 g/L, respectively, and volume fractions of up to 20%^1–4^. This molecular crowding significantly influences cytoplasmic rheology, i.e., how the cytoplasm flows and deforms. In fact, many essential cellular processes and structures, including the assembly of the divisome and the regulation of transcriptional activity by transcription factors, rely on molecules and molecular complexes being able to diffuse within the cell, so that they can encounter and interact with their partners^5–7^. As such, understanding the rheology of the cytoplasm is a matter of significant relevance.

The cytoplasm cannot be regarded as a simple aqueous solution. Studies have shown that in vivo protein diffusion coefficients are one order of magnitude lower than in solution^8–13^ and show a dependence on protein size (*D ∼ r^k^*, where r is the protein radius and *k* < -1) that cannot be explained by the Stokes-Einstein equation, which stipulates that *k* = -1 in simple fluids^11,14–18^. Further, by probing the rheology of the cytoplasm with both endogenous^19–23^ and exogenous^24–33^ supramolecular assemblies, it has been shown that crowding by ribosomes^29^, as well as physicochemical factors, such as pH^27^, can impose constraints on the diffusion of nanometer-sized structures in the cytoplasm.

Intriguingly, the metabolic state has also been shown to affect the rheology of the cytoplasm. For instance, the diffusion of tracer particles is significantly hampered in cells without metabolic activity^16,25,34^. This conclusion was based on measurements in normally growing cells and cells under “extreme” condition, namely starvation or chemically induced energy-deprivation. One explanation for this observation could be an increase in cellular density, which is known to occur upon depletion of metabolic activity^31^. Alternatively, the lower diffusion coefficients could also result from the loss of the hydrotropic effect of ATP^35^. Nevertheless, the mechanistic connection between metabolism and cytoplasmic rheology remains only partially understood. It is unclear what happens in conditions of exponential growth, where cellular density is independent of the carbon and energy source^36,37^.

In this study, we used *E. coli* expressing a 40 nm-sized encapsulin-based fluorescent particle^29^ to probe the rheology of the cytoplasm when cells are growing exponentially on different nutrients, and thus metabolic conditions. We observed a 3-fold variation in the particle’s diffusion coefficient across those conditions. This condition-dependence could not be explained by known determinants previously suggested to affect the rheology of the cytoplasm, such as ribosome abundance, intracellular pH or ATP concentration. Instead, we discovered a robust anticorrelation with the abundance of proteins involved in amino acid metabolism (COG-E). We found that these proteins are more hydrophobic (i.e., have higher hydropathy scores) and more compact than others, and are capable of assembling into agglomerates of folded proteins. Under conditions where COG-E proteins are highly expressed, agglomerates become more abundant and obstruct diffusion of the 40 nm particle. By controlling the abundance of agglomerates, the change of the cellular protein make-up across exponential growth conditions impacts the rheology of the cytoplasm, revealing a previously underappreciated level of spatial organization.

## Results

### Diffusion of 40-nm particles varies with growth conditions

We expressed a genetically-encoded, self-assembling fluorescent particle with a diameter of approximately 40 nm^29^ (PfVS) in the cytoplasm of exponentially growing *Escherichia coli*. The growth medium was supplemented with substrates that require glycolysis or gluconeogenesis, either as single carbon- and energy sources, or with casamino acids, as additional sources of nitrogen and carbon (Fig. 1A). We additionally used a Δ*ptsG* mutant strain, which has a disrupted glucose-specific phosphotransferase system^38^, to measure diffusion under conditions of impaired glucose uptake. We ensured steady-state growth conditions, by propagating cells continuously in the same media for at least eight generations, and by avoiding entry into stationary phase. In addition to these exponential growth conditions, we also analyzed starved cells, i.e., cells that were washed and kept in nutrient-free medium for 7 – 8 hours. Across the twelve exponential growth conditions, specific growth rates (henceforth simply “growth rates”) varied from 0.26 to 1.23 h^-1^ (Fig. 1B), with cell widths from 0.8 to 1.1 µm, and cell lengths from 2.2 to 3.4 µm (Fig. 1C).

**Figure 1.**
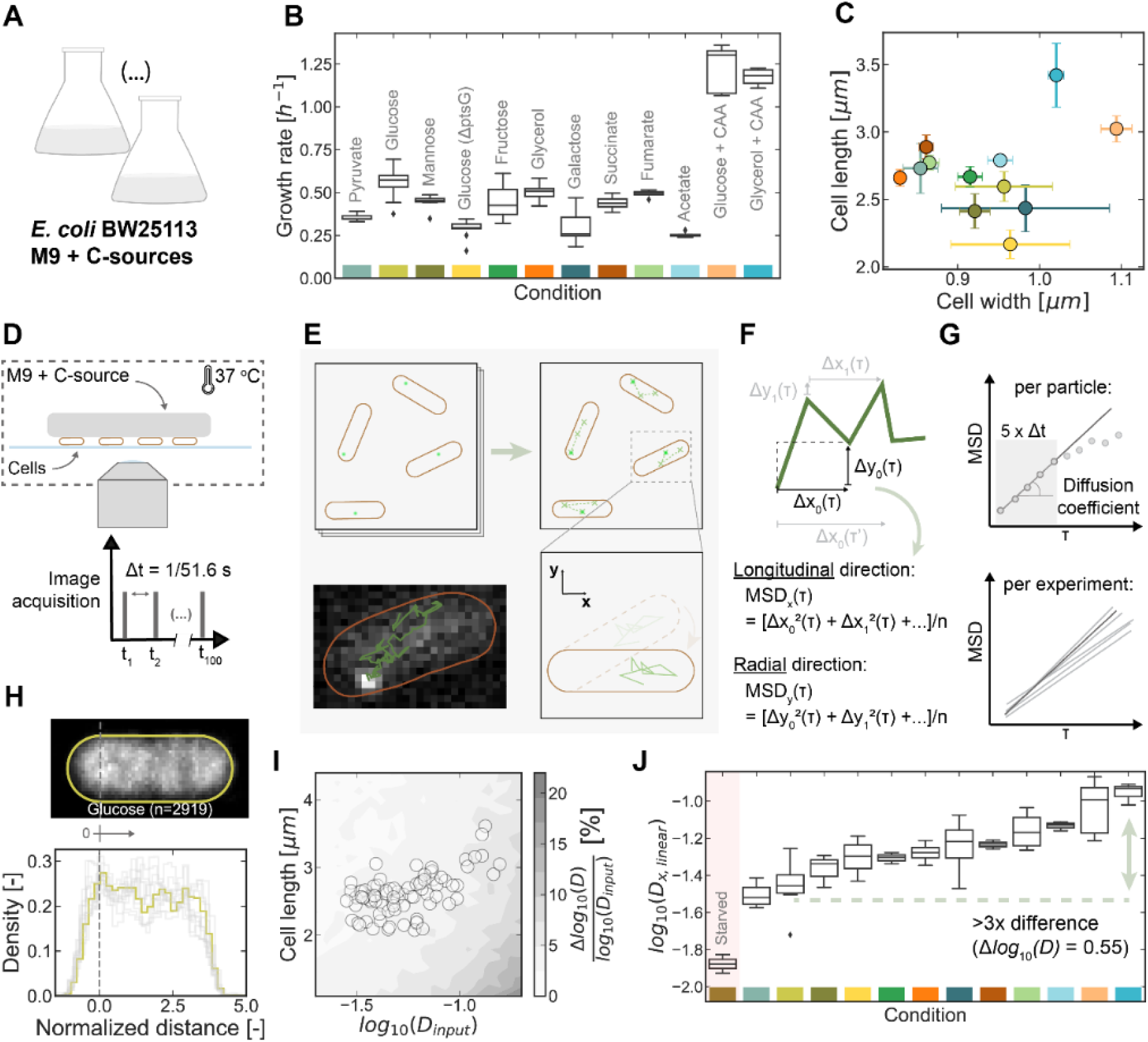
Diffusion of 40-nm particles varies with growth conditions. (A) Schematic of the experimental conditions used. (B) Growth rates for the different conditions of exponential growth. Determined by measuring the cell concentration (cells/mL) as a function of time, prior to sampling for microscopy. Boxes show the quartiles of the distribution across n ≥ 3 experimental replicates. Colors at the bottom of the graph are shown for easier identification of each growth condition. (C) Average cell width and length in different growth conditions. Large variability of cell width during growth on galactose results from the appearance of a subpopulation of cells with different morphology (see Fig. S1B). Cells with visible constriction rings were ignored. Markers show mean ± standard deviation of n ≥ 3 experimental replicates where, for each replicate, the mean of the distribution of single-cell width and single-cell length was used. Same colors as in (**B**). (D) Schematic of the setup used for microscope image acquisition. (E) Schematic of particle tracking and rotation, with an example trajectory (bottom left). (F) Schematic showing the calculation of the mean squared displacement along the long or short cell axes, *MSDx* and *MSDy*, respectively. Exemplified for a time lag of τ=2*Δt. (G) Schematic showing the determination of the diffusion coefficients from the MSD, by fitting the first five time lags to an equation of the form MSD = f(τ, D) (Eq. 1, 2 or 3). (H) Two-dimensional histogram of particle localizations across n = 2919 cells grown on glucose (top), and its one- dimensional projection along the long axis (bottom, colored). Remaining conditions shown in gray. Vertical lines in both panels indicate the origin of the long cell axis. (I) Discrepancy between experimentally determined diffusion coefficients and the confinement-corrected values, obtained through simulations, for a fixed cell width of 0.8 µm and variable length. Markers represents the average cell length and diffusion coefficient measured in individual experiments, under the same conditions as in (**B**). (J) Average (log10) diffusion coefficient along the long cell-axis, across conditions of exponential growth and starvation (shaded). Diffusion coefficients were obtained by fitting MSD plots with Eq. 1 and their original units are µm^2^/s. Boxes show the quartiles of the distribution across n ≥ 3 experimental replicates, except for the “Starved” condition, for which n = 2. See also Figure S1.

To determine the particle’s diffusion coefficients, we transferred samples of exponentially growing cells expressing the fluorescent particle to glass coverslips, covered them with agarose pads containing the same medium as in the culture flask, and acquired images at 51.6 frames per second at constant temperature, 37 °C (Fig. 1D). Analyses of the time-lapse images with a single particle tracking algorithm^39^ detected one or two particle trajectories per cell (Fig. S1A), each trajectory being visible for up to 100 frames. Then, we rotated the cell images and the particle trajectories, so that the x- and y- coordinates describe motion along the long- and short- axes of each cell, respectively (Fig. 1E), and determined the mean squared displacement, *MSD*, of each particle trajectory using those longitudinal- and radial- coordinates (Fig. 1F). Finally, we estimated the diffusion coefficient of each particle, *D*, by fitting the *MSD* values of the first five time lags, corresponding to an interval of <100 ms, to equations describing either normal (*MSD* = 2*Dlinearτ* + *c*, Eq. 1; or *MSD* = 2*Dappτ*, Eq. 2) or anomalous diffusive motion (*MSD* = 2*Γτ^α^*, where *Γ* is the generalized diffusion coefficient; Eq. 3; Fig. 1G).

Importantly, the nucleoid or the cellular membrane could impact our diffusion measurements due to nucleoid exclusion or spatial confinement. However, previous work^32,40^ and our observations of a uniform particle distribution across the whole cell length (Fig. 1H) suggested that 40 nm particles are not excluded from the *E. coli* nucleoid. As cell shape differs markedly across growth conditions (Fig. 1C), we performed computational simulations^41^ of particles moving with Brownian motion in cell-shaped and -sized compartments to assess potential confinement by the cell membrane. As compartment width, we chose the most confining scenario, i.e., a cell width of only 0.8 µm (Fig. 1C), whereas for the compartment lengths and the ground-truth diffusion coefficient of the particles, *Dinput*, we chose the values observed in the various experimental conditions. We then analyzed the simulated particle trajectories with the same approach we used to analyze the experimental particle trajectories (Fig. 1E-G). Using the obtained apparent diffusion coefficients, which potentially include the effect of confinement, we found that the difference between the apparent and the underlying diffusion coefficients in our experimental conditions is, at most, 8% (Fig. 1I). Thus, we conclude that our experimental measurements are not significantly affected by changes in cell shape, i.e., confinement by the cell membrane. This is consistent with our finding that radial and longitudinal diffusion coefficients are indistinguishable (*p* ≥ 0.16, Kruskal-Wallis H-test) in the two cell subpopulations with largely different cell sizes that appear during growth on galactose (Fig. S1B).

Next, focusing on the particle’s longitudinal diffusion coefficients, estimated with a linear model (Eq. ^1^; used henceforth in this work), *Dx,linear*, we compared the results obtained for the different growth conditions. Here, we found that the diffusion coefficient changed by more than 3-fold (i.e., more than 0.48 on a log10-scale) across the range of exponential growth conditions (Fig. 1J). Specifically, the lowest particle mobility was observed on pyruvate (log10(*Dx,linear*) = -1.51 ± 0.06) and the highest on glycerol supplemented with casamino acids (log10(*Dx,linear*) = -0.95 ± 0.06). The condition-dependence of the diffusion coefficients was robust to the use of different data processing steps (Fig. S1C) and diffusion models (Eq. 2 and 3; Fig. S1D). Furthermore, consistent with previous observations of decreased particle mobility in cells with inhibited or significantly reduced metabolic activity^25,34^, we also found lower diffusion coefficients in starved cells than in any of the exponential growth conditions (log10(*Dx,linear*) = -1.88 ± 0.07). Overall, our findings unexpectedly show that particle mobility in the *E. coli* cytoplasm varies between conditions of exponential growth approximately as much as between exponential growth and starved conditions. This finding raises the question of what causes such large differences.

### Diffusion coefficients correlate with COG-E protein abundance

One parameter that could affect the diffusion coefficient of the particle across the conditions is cell density, i.e., the mass of dry material per unit of cell volume^36^. However, with various techniques, it was found that this parameter remains essentially unchanged when *E. coli* exponentially grows on different nutrients^36,37,42–44^. Thus, cell density is unlikely to explain the 3-fold difference between the diffusion coefficients we measured across conditions of exponential growth.

We next considered previously proposed determinants of cytoplasmic rheology, such as ribosome abundance^29^, intracellular pH^27^ and ATP concentration^35^. First, ribosome abundance is known to scale approximately linearly with growth rate^45–47^. Thus, if ribosomes hindered the diffusion of the 40 nm particle, we would expect an anticorrelation between the diffusion coefficient and growth rate. Instead, if at all, our data shows the opposite trend (Fig. 2A), rendering ribosome abundance an unlikely explanation for the condition-dependent changes of diffusion. Second, having measured intracellular pH with a pH sensor, pHluorin^48^, we also did not find any correlation between the diffusion coefficients and intracellular pH (Fig. 2B). Furthermore, by combining diffusion and pH- measurements in cells that were exposed to the metabolic inhibitor 2-deoxyglucose, we found that significant drops in intracellular pH lead to only minor decreases in the diffusion coefficients (Fig. S2A). These findings suggest that the observed differences in diffusion cannot be explained by pH either. Third, we did not find any correlation between the diffusion coefficients and intracellular ATP concentrations reported for the different growth conditions^49–51^. These findings suggest that likewise ATP – a hydrotrope that could fluidize the cytoplasm^35^ – cannot explain the changes in the diffusion of the 40 nm particle across the exponential growth conditions (Fig. 2C and Fig. S2B-D).

**Figure 2.**
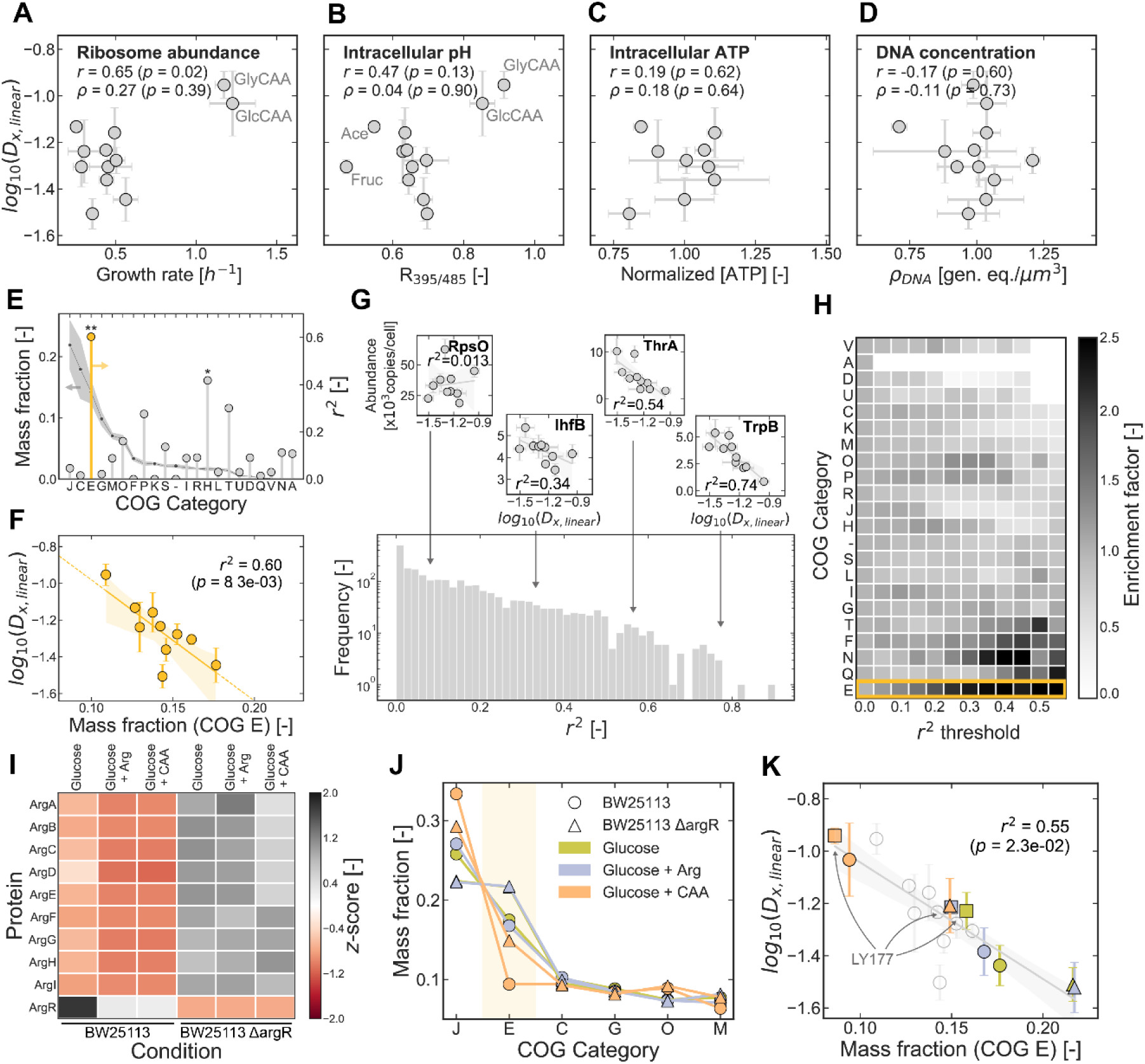
**Diffusion coefficients correlate with COG-E protein abundance** (A) Average diffusion coefficients of all exponential growth conditions shown in Fig. 1J plotted against growth rate, as a proxy for ribosome abundance. Pearson (*r*) and Spearman (*ρ*) correlation coefficients are displayed. (B) Similar to (**A**) using the fluorescence readout of pHluorin, as a proxy for intracellular pH. Outliers are indicated: "Ace", acetate; "Fruc", fructose; "GlyCAA", glycerol and casamino acids; "GlcCAA", glucose and casamino acids. (C) Similar to (**A**) for the normalized intracellular ATP concentration, as reported by Radoš *et al*^51^. See Fig. S2B-D for color-labeled conditions and other metabolomics data sets. (D) Similar to (**A**), for the volumetric concentration of DNA, *ρDNA*, estimated based on the average cell volume under each condition and estimated number of genome equivalents, reported by Bremer & Dennis^62^. (E) Average mass fraction of each COG category across growth conditions (continuous line; left vertical axis), as reported in Schmidt *et al*^54^’s data set. Values of the squared Pearson correlation coefficient, *r^2^*, between diffusion coefficients and protein mass fraction are shown on the right vertical axis (stems). “*”: *p* < 0.05, “**”: *p* < 0.01 with a two-sided Wald-test. (F) Correlation between the measured diffusion coefficients and mass fraction of COG category "E" ("Amino acid transport and metabolism"), highlighted in (**E**). (G) Histogram of *r^2^* values obtained when the absolute abundances, in copies/cell, of all 2357 proteins in the proteomics data set (Schmidt *et al*^54^) are plotted against the diffusion coefficients measured under the same growth conditions, shown for four example proteins. Note the logarithmic scale on the y-axis. (H) Fractional enrichment of COG categories among the set of proteins with high correlation coefficients. The enrichment factor is the ratio between the fraction of proteins that belong to a COG category and have *r^2^* ≥ *r^2^*, and the fraction of all proteins in the proteome that belong to that COG category. (I) Normalized abundance (z-score) of ArgR and other nine proteins under its regulation, in experimental conditions designed to alter the abundance of amino acid metabolism proteins. (J) Mass fraction of the six most abundant COG categories under the new experimental conditions. J, "Translation, ribosomal structure and biogenesis"; C, "Energy production and conversion"; E, "Amino acid transport and metabolism"; G, "Carbohydrate transport and metabolism"; O, "Posttranslational modification, protein turnover, chaperones"; and M, "Cell wall/membrane/envelope biogenesis". Values for the Δ*argR* strain on glucose and glucose + arginine overlap. (K) Mass fraction of COG-E proteins plotted against the diffusion coefficients measured under the same conditions. White circles: BW25113 strain in the original set of conditions (same as in (**F**)); squares: LY177 strain^52^; other symbols and colors as indicated in (**J**). Grey line was obtained by linear regression of the entire data set. Shaded area is the 95% confidence interval of the regressed line, obtained by bootstrapping. Colored markers show the mean ± standard deviation of n ≥ 3 experimental replicates. Original units of the diffusion coefficients are µm^2^/s. See also Figure S2.

Finally, we asked whether changes in the size of the nucleoid, with its mesh-like structure, could be responsible for the condition-dependent diffusion coefficients. However, neither the estimated DNA concentration (Fig. 2D) nor the nucleoid-to-cytosol ratio^30^ (Fig. S2E) varied monotonically with the measured diffusion coefficients, leading us to also exclude DNA as a parameter to explain the observed trend. Thus, none of the parameters previously suggested to impact the diffusion of similarly sized particles could explain the condition-dependence of the diffusion coefficients we observed.

Yet, as we have recently found that proteins and ribosomes were responsible for most of the hindrance experienced by the 40 nm particle in vivo^52^, and because it is known that *E. coli* adapts its proteome when grown on different carbon sources^46,53–55^, we wondered if certain proteins could be responsible for the observed condition-dependence of the diffusion coefficients. We first asked whether the abundance of any subset of functionally related proteins would correlate with the measured diffusion coefficients. We used the proteomics data of Schmidt *et al*^54^, determined the mass fraction of each functional category of proteins (i.e., “Clusters of Orthologous Groups”, COG^56^) in the ten growth conditions for which we had measured diffusion coefficients, and then computed the correlation between these mass fractions and diffusion coefficients. Here, out of the 22 COG categories, we found that only category E, corresponding to “Amino acid transport and metabolism”, showed a high and statistically significant correlation (*r^2^* = 0.60; *p* = 0.008; Fig. 2E-F). Notably, this COG category is the third most abundant category of proteins in *E. coli* by mass, with its mass fraction ranging between 11% and 18% across the investigated conditions (Fig. 2F). Thus, a high protein abundance of the COG-E category correlates with low particle mobility.

As this anti-correlation could be caused by solely a few, highly abundant proteins of that category, as opposed to being a general trend of COG-E proteins, we repeated the correlation analysis using the abundances of all 2357 proteins in the proteomics data set, one at a time. While most proteins correlate weakly with the diffusion coefficients (such as RpsO or IhfB), we found that 4.5% of all 2357 proteins had correlation coefficients higher than *r^2^* ≥ 0.5 (such as ThrA and TrpB; Fig. 2G). An enrichment analysis revealed that proteins of the COG category “E” were over-represented among strongly correlating proteins (Fig. 2H and Fig. S2F-G). Thus, more than proteins of any other category, proteins involved in amino acid metabolism (i.e., COG-E proteins) display a robust anticorrelation with diffusion coefficient of the 40 nm particle.

We then asked whether the observed trend of the diffusion coefficients would generalize to additional conditions, in which we perturbed the mass fraction of COG-E proteins. To this end, we used a Δ*argR* knockout strain and grew these cells in medium supplemented with amino acids. Since *argR* encodes a transcription factor that regulates the expression of genes involved in the arginine biosynthesis pathway^57–59^, *argR* deletion results in increased abundance of the enzymes of this pathway^60^. We supplemented the culture medium with either L-arginine or casamino acids to further modulate the abundance of these proteins. When these amino acids are present in the medium, the expression of the relevant biosynthetic enzymes is downregulated^53,54,60,61^. We confirmed through proteomic analyses that the enzymes involved in arginine biosynthesis are indeed expressed more in Δ*argR*, and expressed less in the wild-type strain in the presence of externally supplied amino acids (Fig. 2I). With these experimental and genetic perturbations, we successfully altered the mass fraction of COG-E proteins (Fig. 2J). The diffusion coefficients measured under these new conditions, plotted versus the corresponding mass fractions of COG-E proteins, maintained the same anticorrelation trendline (Fig. 2K).

To further probe the robustness of this anticorrelation, we turned to another *E. coli* strain, LY177, for which we had previously found diffusion coefficients that were systematically higher (Δlog10(*D*) ≥ 0.09) than for the BW25113 strain used here^52^. If the abundance of COG-E proteins has a causal role in determining the diffusion coefficient, we hypothesized that this strain would have a lower abundance of COG-E proteins. Indeed, proteomic analyses revealed a lower abundance of COG-E proteins (16% vs 18% for LY177 and BW25113 on glucose, respectively), likewise preserving the anticorrelation with the diffusion coefficients (Fig. 2K). Notably, we saw that in conditions where growth rates are comparable, but diffusion coefficients are different, the abundance of COG-E proteins was also different (Fig. S2H-J), suggesting ribosome abundance was not the reason for the condition-dependent changes in diffusion. Finally, we also excluded mRNA encoding for COG-E proteins (Fig. S2K-L), as well as intracellular amino acid concentration (Fig. S2M), as responsible for the observed correlations.

Together, these findings show that there is a strong anticorrelation between the abundance of COG- E proteins and the diffusion coefficient of the 40 nm particle. This correlation is retained upon various types of perturbations of the protein abundances, suggesting a causal relationship.

### COG-E protein abundance increases cytoplasmic viscoelasticity

To investigate how COG-E proteins could slow down the diffusion of the 40 nm particle, we considered two scenarios. First, COG-E proteins could display stronger intermolecular interactions, which could make the cytoplasm more viscous and slow down the diffusion of the particle. Second, COG-E proteins could form larger obstacles with which the particle collides, turning the cytoplasm into a viscoelastic environment (Fig. 3A). Seeking to distinguish between these two scenarios, we first assessed if the diffusive behavior of the particle reveals viscoelasticity. For this, we calculated the velocity autocorrelation function using the particle trajectories collected under all conditions in our dataset. The velocity autocorrelation function, 𝐶^𝛿^/𝐶^𝛿^(𝜏 = 0), measures the alignment between displacements undergone by each particle during a period *δ*, and separated in time by a lag *τ* (Fig. 3B). Importantly, when *τ* = *δ*, this function measures the alignment between *consecutive* displacements. We observed that the autocorrelation function consistently displayed negative values when *τ* = *δ* over a broad range of timescales (Fig. 3C), revealing that consecutive particle displacements are negatively correlated. This means that the diffusion of 40 nm particles is non- Brownian and there is a degree of “push back” as they move through the cytoplasm, suggesting that large obstacles exist in their paths and indicating viscoelasticity^63^ (Fig. 3B).

**Figure 3.**
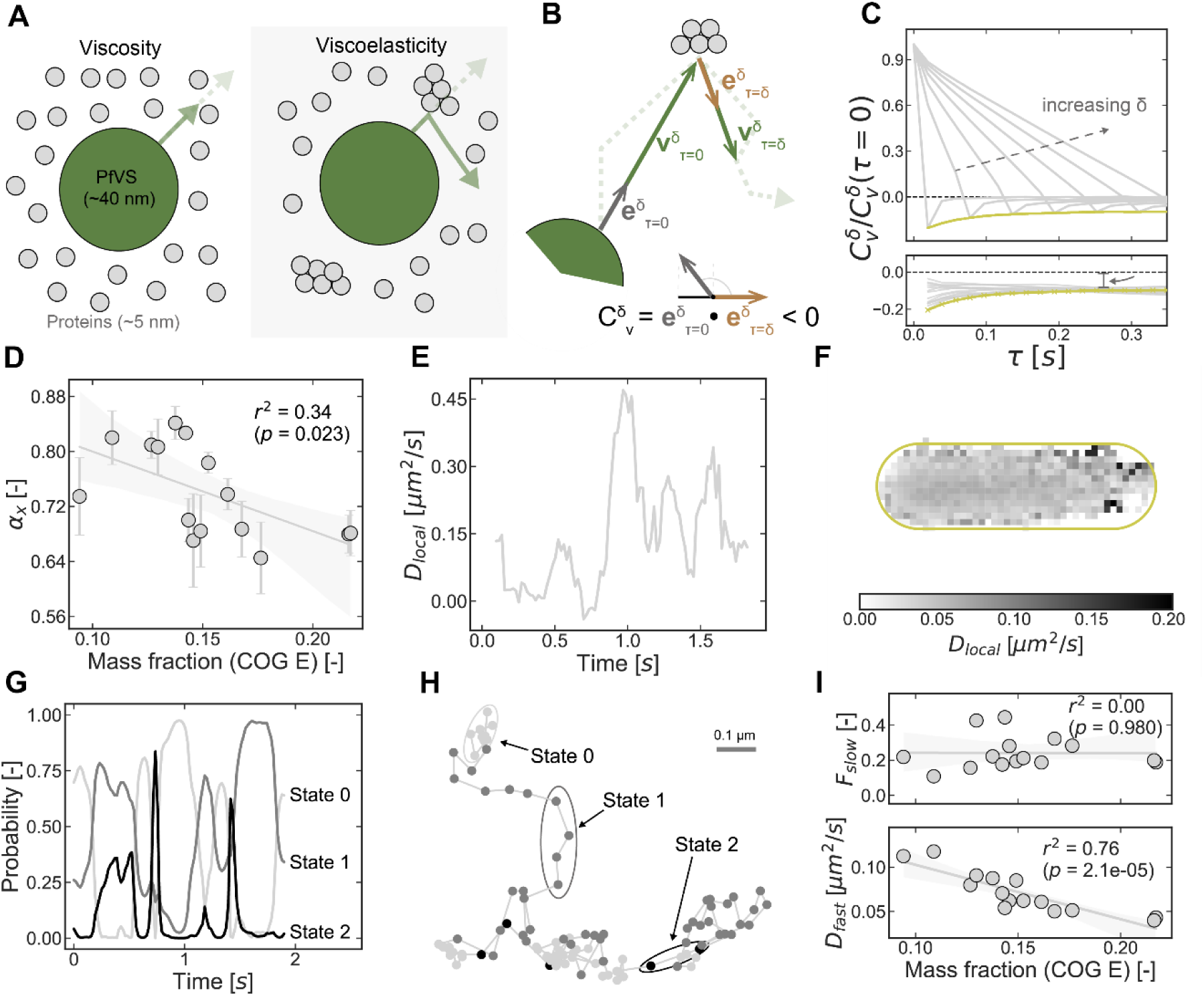
**COG-E abundance increases cytoplasmic viscoelasticity.** (A) Schematic of two scenarios for how cytoplasmic proteins (grey circles) could affect the diffusion of the 40 nm particle (green circle). (B) Schematic showing how the velocity autocorrelation function is determined, as the dot product of two unit vectors, ***e***, representing the direction of the displacement undergone by the particle between *τ* and *τ + δ*. In this example, *τ = δ* = 2 time intervals. The “pushback” of the particle towards its original position results in a negative value for their dot product, 𝐶^𝛿^. This calculation is repeated for the ensemble of all particles, and across all time points. (C) Normalized velocity autocorrelation function, VAF, for the glucose growth condition, as a function of the time lag, *τ,* and for increasingly large periods, *δ* (increasing from left to right; upper panel). Colored line traces the negative “peaks” of the VAF across different values of *δ*. Peak profiles of VAF curves obtained for glucose (yellow) and remaining growth conditions (grey) are shown on the lower panel. Arrow indicates that these profiles remain below zero for all time lags, a hallmark of viscoelasticity. (D) Anomalous diffusion exponent, *αx*, for each growth condition as a function of the mass fraction of COG-E proteins. Values of *αx* were obtained by fitting the longitudinal *MSD* (“x”) of each particle to Eq. 3. (E) Heterogeneity of diffusive behavior of a single particle trajectory, assessed by its local diffusion coefficient as a function of time. Values of *Dlocal*, were determined from the *MSD* in a sliding time window of ten timesteps. (F) Map of the average *Dlocal* across the cell, obtained by overlaying the *Dlocal* for all trajectories detected on the glucose growth condition and by averaging in spatial bins. (G) Probability of an example particle being in any of three diffusive states (states 0, 1 or 2) as a function of time. State properties (diffusion coefficients and fractions) and local state probabilities were obtained with *ExTrack*. (H) Trajectory of the particle analyzed in (**G**), annotated with the color of the most likely diffusive state. (I) Fraction of the slowest diffusive state (i.e., state 0; upper panel) and average diffusion coefficient of the remaining two states (lower panel), plotted against the mass fraction of COG-E. Fractions and diffusion coefficients of states 1 and 2 shown in Fig. S3B. Line obtained by linear regression. Shaded area is the 95% confidence interval, obtained by bootstrapping. See also Figure S3.

Another hallmark of diffusion in a viscoelastic environment is that particles display a “subdiffusive” type of motion^64,65^. For each growth condition, we thus determined the anomalous diffusion exponent, *α*, which describes how the *MSD* scales with *τ* (see Eq. 3) and quantifies how a particle’s diffusive behavior differs from that in a purely viscous environment, for which *α* is 1. We observed that, akin to the diffusion coefficients, *α* decreases with the increasing abundance of COG-E proteins (Fig. 3D), showing that particles not only diffuse slower, but also in an increasingly subdiffusive manner. Together, our findings show that the particles experience the cytoplasm as an increasingly viscoelastic environment as the abundance of COG-E proteins increases. This result suggests that COG-E proteins could be forming large assemblies in the cytoplasm.

Given that COG-E proteins account for only about 11-18% of the proteome^54^, we reasoned that if they indeed form large assemblies, these structures should be sparsely and heterogeneously distributed in the cytoplasm. As a result, we expected the diffusion of the 40 nm particle to be heterogeneous, as the particle occasionally encounters such assemblies. Visual inspection of individual tracks indeed suggested changes of diffusivity along time. Then, we determined local effective diffusion coefficients, *Dlocal*, on a sliding window of ten timesteps along the trajectories (exemplified in Fig. 3E). We found high variations of *Dlocal* through time that cannot be explained by free diffusion in a homogenous environment (Fig. S3A). We also found that low values of *Dlocal* are not linked to cellular regions such as the pole or the nucleoid (Fig. 3F). Reinforcing that the diffusion of the particle is highly heterogeneous, using a probabilistic model, *ExTrack*^66^, we could identify at least three different diffusive states for each growth condition: a state of nearly immobile particles (state 0) and two states of faster diffusion (states 1 and 2; Fig. 3G-H). While the fraction of nearly immobile particles (state 0) was independent of the abundance of COG-E proteins (Fig. 3I, top), we found that the combined diffusion coefficient of the faster states anticorrelates with the abundance of COG-E proteins (Fig 3I, bottom and Fig. S3B).

Together, our data show that (i) the diffusion of the 40 nm particle is reduced and (ii) the cytoplasm becomes increasingly viscoelastic as the abundance of COG-E proteins increases, with (iii) the particle showing heterogeneous diffusion. We therefore hypothesized that the heterogeneity and viscoelasticity result from the formation of protein structures large enough to obstruct the movement of the 40 nm particle.

### In vivo formation of protein agglomerates

To investigate whether COG-E proteins form supramolecular assemblies in vivo, we examined cells using transmission electron microscopy (TEM). We specifically looked for one type of such assemblies – several nanometer-long protein filaments – which has been observed in bacteria under certain conditions^67–69^. To this end, we imaged cells grown on glucose, glycerol and fumarate, reflecting different diffusion coefficients of the 40 nm particle (Fig. 1J), and presumably variable abundance of protein assemblies. However, we could not find any such filamentous structures (Fig. 4A). Thus, if COG-E proteins form supramolecular assemblies, these must either be smaller than the previously described filaments (up to a few dozen nm) or indiscernible in TEM images of a crowded cytoplasm.

**Figure 4.**
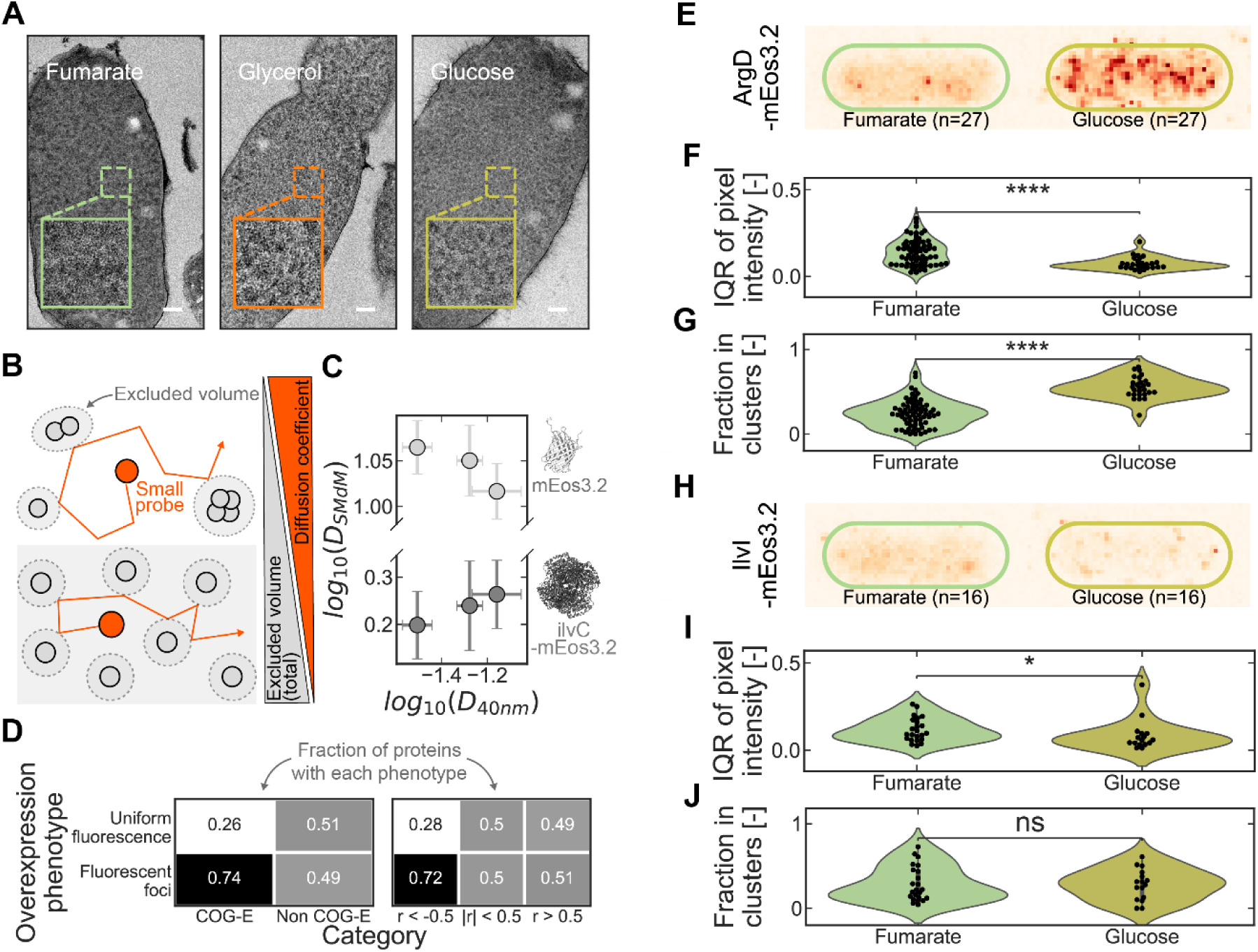
In vivo formation of protein agglomerates. (A) Transmission electron microscopy images of cells grown on fumarate, glycerol, or glucose. The white patches visible on all three images are artifacts. Insets are close-ups of the central regions. Scale bar: 100 nm. (B) Schematic showing how the formation of protein assemblies (grey, top) could lead to increased mobility of protein- sized fluorescent probes (dark orange) relative to conditions where the same proteins remain separate (grey, bottom). Steric interactions reduce the space available to the fluorescent probe. (C) Diffusion coefficients of two fluorescent proteins, mEos3.2 (monomer) and ilvC-mEos3.2 (homotetramer), obtained by single molecule displacement mapping in medium with either pyruvate, glycerol or fumarate, *DSMdM*. These values are plotted against the diffusion coefficient of the 40 nm particle, *D40nm*, in the same growth conditions. Molecular structures of both proteins, obtained with *AlphaFold Multimer,* are shown next to the corresponding data points. Values of the SMdM diffusion coefficients are mean ± standard deviation of n ≥ 20 cells. Original units of the diffusion coefficients are µm^2^/s. (D) Fractions of proteins that are uniformly distributed or form fluorescent foci in the cytoplasm, when overexpressed as GFP-fusions in *E. coli,* according to Györkei *et al*^71^. Values add up to 1 along each column. The total numbers of proteins per category are: *nCOG-E* = 145; *nNon COG-E* = 1232; *nr<-0.5* = 127; *n|r|<0.5* = 1067; *nr>0.5* = 183. Test for equivalence in the proportions of overexpression phenotypes in the different groups: *p* = 2.0 x 10^-8^ (COG-E vs Non COG-E); *p* = 2.6 x 10^-6^ (*r* < -0.5 vs *r* ≥ -0.5), with Fisher’s exact test (two-sided). (E) Two-dimensional histograms of protein blinking events in cells expressing ArgD-mEos3.2, fixed with formaldehyde after growing on fumarate (left) or glucose (right). Same number of randomly drawn cells used to generate each histogram, *n*, was chosen for a fair comparison of the heterogeneity. For this, we *undersampled* the condition with the highest number of cells imaged. For example: we imaged 67 cells expressing ArgD-mEos3.2 grown on fumarate, but only 27 on glucose; thus, we randomly selected 27 of the cells grown on fumarate. (F) Distribution of *IQR* values of the normalized pixel intensities for cells expressing ArgD-mEos3.2. Each black marker represents a single cell. (G) Similar to (**F**), showing the fraction of clustered blinking events. “ns”: non-significant, “*”: 0.01 < *p* < 0.05, “****”: *p* < 1 x 10^-4^ with a Mann-Whitney U-test. (**H-J**) Same as (**E-G**) for cells expressing IlvI-mEos3.2. See also Figure S4.

We then sought to estimate the size of such assemblies via indirect measurements. Here, we reasoned that the local increase of protein concentration arising from the formation of structures should free up space elsewhere in the cytoplasm. Any tracer particle that is sufficiently small to experience this increase of “free space” should therefore diffuse faster^70^ (Fig. 4B). Based on this reasoning, we anticipated that diffusion measurements with differently sized particles should provide an indication of the size of the obstacles in the cytoplasm. Accordingly, we used single molecule-displacement mapping (SMdM) to determine diffusion coefficients of two differently sized proteins, monomeric mEos3.2 (26 kDa, ∼5 nm in diameter) and homotetrameric IlvC-mEos3.2 (319 kDa; ∼10 nm in diameter) in three growth conditions for which we had observed different diffusion coefficients of the 40 nm particle (i.e., medium with pyruvate, glycerol and fumarate; Fig. 1J). Akin to the 40 nm particle, we found that the larger IlvC-mEos3.2 diffused more slowly on pyruvate and glycerol than on fumarate, whereas the smaller mEos3.2 displayed the reverse trend (Fig. 4C). As the same growth conditions affect the diffusion of these two differently-sized fluorescent proteins in opposite directions, this suggests that the “free space” changes across growth conditions (Fig. 4B), and that this is already perceived on the length scale of ∼5 – 10 nm.

To further investigate the existence of protein assemblies, we turned to a recent study showing that certain *E. coli* proteins, when fused to GFP, could form “foci” rather than distributing uniformly throughout the cytoplasm^71^. We wondered if COG-E proteins would display a tendency to form such foci. We analyzed the dataset of Györkei *et al*^71^ (Table S1 therein), assigned COG categories to each protein and performed an enrichment analysis. Here, we found that COG category E was enriched in proteins which form foci (*p* = 2.0 x 10^-8^, Fisher’s exact test Fig. 4D, left). Likewise, we found that the group of “anticorrelating” proteins (i.e., proteins whose abundances correlate with diffusion with *r* < -0.5) was enriched in proteins that form fluorescent foci (*p* = 2.6 x 10^-6^, Fisher’s exact test; Fig. 4D, right). Thus, many of the proteins potentially implied in the slowdown of diffusion of the 40 nm particle tend to form supramolecular assemblies.

Since Györkei *et al*’s study relied on protein overexpression, we asked whether native-level expression of COG-E proteins would similarly lead to clustered localization, and whether any clustered localization was condition-dependent, as expected from the condition-dependent viscoelasticity of the cytoplasm (Fig. 3C-D). We thus tagged two COG-E proteins, ArgD and IlvI, with the photoswitchable protein mEos3.2, and expressed these fusion proteins from their native chromosomal loci and promoters. We grew cells on either glucose or fumarate, fixed them with formaldehyde and used photoactivated localization microscopy (PALM) to obtain super-resolution images. For ArgD, we observed a more heterogeneous distribution of fluorescence on glucose than on fumarate, with multiple clusters distributed throughout the cytoplasm (Fig. 4E). Quantification of the fluorescence heterogeneity in these cells using the interquartile range (IQR) of the normalized pixel intensity distributions revealed higher heterogeneity, i.e., a few bright pixels and a lower IQR, on glucose than on fumarate (Fig. 4F and S4A). Notably, glucose is the condition in which COG-E proteins are more abundant and in which we expected more clusters. We arrived at the same conclusion when the heterogenous distribution of fluorophore detections, i.e., “blinking events”, in the PALM images was analyzed with a clustering algorithm (Fig. 4G and S4B). For IlvI, the results were qualitatively similar, although with lower statistical significance (Fig. 4H-J). Thus, COG-E proteins can indeed form supramolecular assemblies in vivo at native expression levels, and the formation of these assemblies is condition-dependent, as we also expected from observing the changing viscoelasticity of the cytoplasm. The appearance of these assemblies throughout the cytoplasm is consistent with the varied local diffusivities (Fig. 3E).

### Protein hydrophobicity and compactness likely drive agglomeration

Next, we aimed to understand how COG-E proteins form assemblies that impede the diffusion of the 40 nm particle. To do this, we sought to identify structure- or sequence-based features that differentiate anticorrelating proteins from those whose abundance does not correlate with diffusion. We trained a random forest classifier to distinguish anticorrelating (*r* < -0.5) from non-correlating proteins (|*r*| < 0.3). This model (Model #1) was provided with 76 structure- and sequence-based features of each protein, derived from various computational tools^72–77^ (Table S1; Fig. 5A). We ensured that strongly collinear features were removed prior to training. Reasoning that only proteins expressed at high copy numbers can affect the diffusion of the 40 nm particle, we restricted our analysis to proteins with an average abundance across the growth conditions of >100 copies/cell. After training, we evaluated the model’s performance on a test set of unseen proteins. This model achieved an area under the receiver operating characteristic curve (AUC-ROC) of 0.72, outperforming a random classifier model, with an expected AUC of 0.5 (Fig. 5B, left). By analyzing the permutation importances of each feature, we found that the “disorder score” was most important for distinguishing between the two protein classes (Fig. 5B, right). Notably, anticorrelating proteins had lower disorder scores than non-correlating proteins (Fig. 5C). Thus, if there is a high abundance of low disorder proteins, diffusion is low.

**Figure 5.**
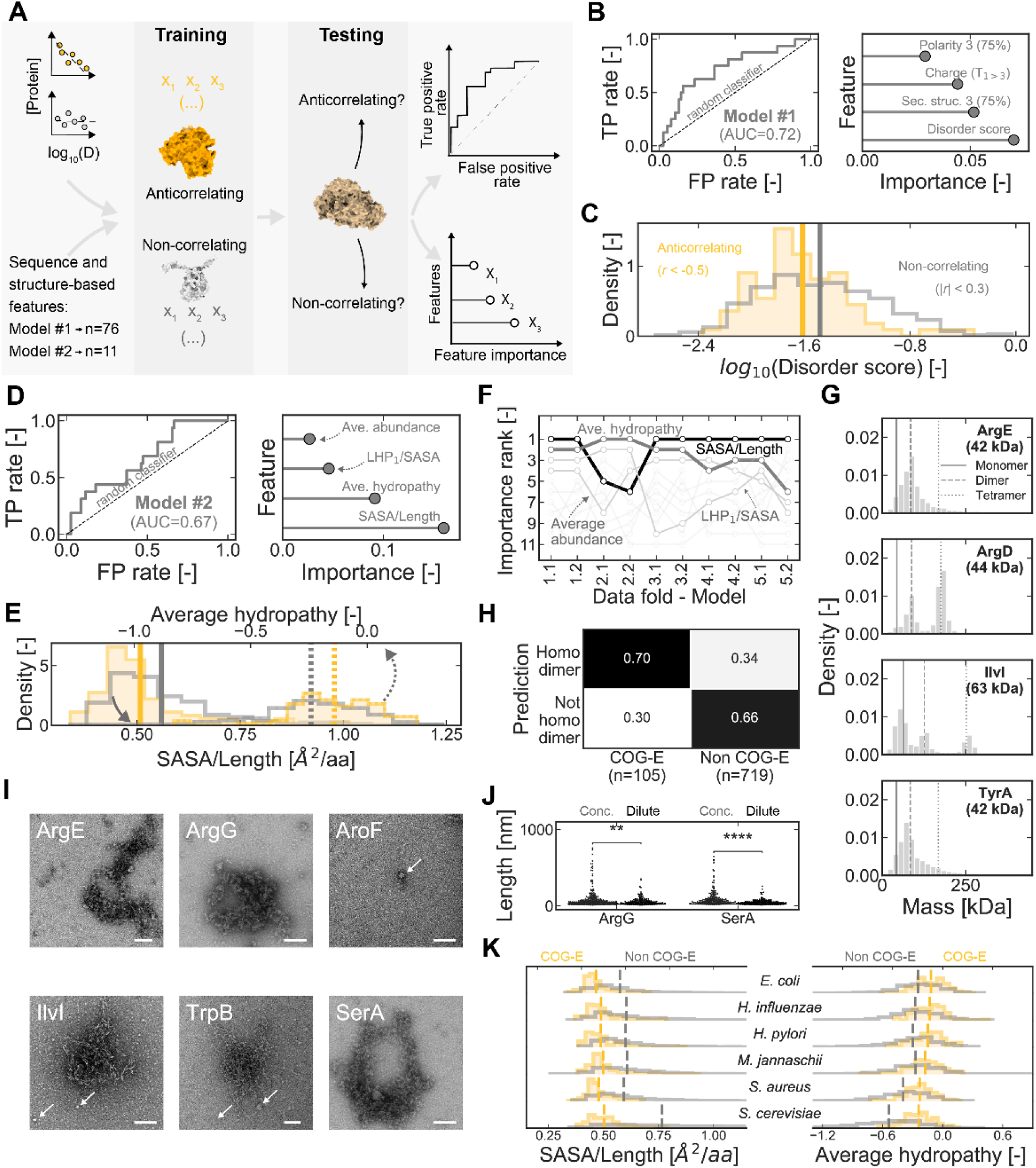
Protein hydrophobicity and compactness likely drive agglomeration. (A) Schematic of the machine learning approach used. Random forest classifiers were trained on sets of either 76 or 11 non-collinear features (*x1, x2, x3*, …) describing each anticorrelating (yellow) or non-correlating protein (grey). The classifiers were tested against proteins unseen during training (brown), and their performance assessed through a receiver operating characteristic curve, ROC (right, top). For the best performing models, we determined the permutation importance of each feature (right, bottom). (B) ROC curve of a random forest model distinguishing anticorrelating (*r* < -0.5) from non-correlating (|*r*| < 0.3) proteins with an average abundance of >100 copies/cell (left). A perfect classifier model would display a ROC curve that passes through the top-left corner of the plot; a random classifier would display a ROC curve that lies on the diagonal. Axes labels: “FP”, false positive; “TP”, true positive. (right) Four most important features of Model #1, ranked by their permutation importances. (C) Distribution of disorder score (determined with *NetSurfP3.0*) in anticorrelating and non-correlating proteins, in log10-scale. Vertical lines indicate the mean of each distribution. (D) Same as (**B**) for Model #2, trained on 11 protein features. (E) Distributions of protein features SASA/L (continuous lines, values shown on the bottom axis) and average hydropathy, according to the Kyte-Doolittle scale, across all amino acids in the protein sequence (dashed lines, values shown on the top axis). Yellow distributions: anticorrelating proteins. Grey distributions: non-correlating proteins. (F) Eleven features ranked according to their permutation importances in models that were trained and tested on different subsets of the data (“data folds”). For every data fold, the two best performing models were selected. A rank of 1 means that the feature had the highest permutation importance. (G) Fraction of cytoplasmic proteins predicted to form homodimers among COG-E and remaining categories. “Homodimers” are proteins with high confidence *AlphaFold Multimer* predictions. Values shown are not explained by random sampling of the entire set of 824 analyzed proteins (*p* = 1.5 x 10^-11^, with a two-sided Fisher’s exact test). (H) Distribution of oligomeric states in samples of purified proteins, obtained by mass photometry. Samples contained dithiothreitol to prevent the oxidation of cysteine residues. The expected mass of the monomer, (homo)dimer and (homo)tetramer are indicated by the vertical lines. (I) Transmission electron microscopy images of purified protein samples, negatively stained with uranyl acetate. Circular structures marked with arrows. Scale bar: 100 nm. (J) Distribution of sizes of irregular-shaped protein agglomerates found in samples of purified ArgG and SerA. The samples were analyzed by EM at either high (“Conc.”) or low (“Dilute”) protein concentrations. For ArgG, the two concentrations used were 0.06 mg/mL and 0.024 mg/mL, and for SerA, 0.06 mg/mL and 0.015 mg/mL. “**”: 1x10^-3^ < *p* < 0.01; “****”: *p* < 1x10^-4^, with Levene’s test centered on the median. Same as (**E**) for six organisms. See also Figure S5 and Tables S1-S3.

As this disorder score is based on non-resolved regions in experimentally determined protein structures^75^, we suspected that the low disorder of anticorrelating proteins would indicate that these proteins have compact and well-defined three-dimensional structures, potentially resulting from a high hydrophobicity. To test this, we selected a set of 17 different features describing hydrophobicity and three-dimensional shape, of which some were not used in the first model (Table S2). After removing collinear features, we trained another random forest classifier (Model #2) using the 11 remaining features. This new model had a performance similar to that of Model #1 when evaluated against an unseen test set (AUC = 0.67; Fig. 5D, left). The most important features of this model were the average solvent-accessible surface area per residue (SASA/L, permutation importance = 0.16) and the average hydropathy score (permutation importance = 0.09; Fig. 5D, right). Consistent with our expectation, anticorrelating proteins have lower values of SASA/L and higher values of average hydropathy (Fig. 5E). We repeated the random splitting of the original data into training and test sets four additional times, and consistently identified SASA/L and average hydropathy as top ranked features (Fig. 5F), showing that the identification of these two features is robust to changes in the input data. Thus, those proteins which are expressed at high levels and are overall more hydrophobic and compact than others are more likely to display an anticorrelation with the diffusion coefficient of the 40 nm particle.

We suspected that proteins with such features could form amorphous assemblies of folded proteins, so-called “agglomerates”^78^. The formation of this type of assemblies occurs when proteins can establish multivalent interactions, i.e., interact via multiple surface regions. As the appearance of symmetrical structures that follows from homo-oligomerization can amplify multivalency, even small assemblies like homodimers could have an enhanced propensity to form agglomerates. To test whether proteins for which we had found a strong anticorrelation with the particle’s diffusion coefficient can indeed form agglomerates, we expressed and purified twelve such proteins. First, we used mass photometry, a technique to determine the mass distribution of macromolecules in solution^79^. We observed homooligomers in all samples, with all showing at least homodimers, and eight proteins forming assemblies of higher order than earlier described (Fig. 5G and S5A). Accordingly, we estimated with *AlphaFold Multimer*^80^ predictions that, among cytoplasmic proteins, approximately 70% of COG-E proteins may form at least homodimers, compared to 34% in the remaining proteome (Fig. 5H). Since mass photometry requires protein concentrations below 50 nM - much lower than the concentration of these proteins in cells (Table S3) - the observed oligomerization suggests that these proteins have high self-binding affinities. Interestingly, we also found that four proteins (ArgE, IlvH, IlvN and TyrA) had broad mass distributions, spanning until >200 kDa, and without clear stoichiometries (Fig. 5G and S5A). This observation suggests the occurrence of large assemblies of undefined oligomeric state, consistent with our hypothesis of agglomerate formation.

To test whether the proteins could indeed form amorphous agglomerates, we performed negative staining EM analysis of purified proteins at micromolar concentrations, resembling reported in vivo concentrations (Table S3). Here, we observed the presence of large sphere- and mesh-shaped structures, occasionally co-occurring in the same sample (Fig. 5I). While spherical structures were found with diameters of up to ∼40 nm, some mesh-like structures could have characteristic lengths of ∼300 nm or more (Fig. 5I). Interestingly, the latter appeared to form reversibly and in a concentration-dependent manner. In samples that were diluted prior to imaging, the structures we found were smaller (Fig. 5J and S5B-C). Although it is unlikely that agglomerates of several hundreds of nanometers in size exist in cells (see Fig. 4A), these findings suggest that proteins which anticorrelate with the diffusion coefficient of the 40 nm particle tend to form more amorphous agglomerates than other proteins of the *E. coli* proteome.

Finally, we asked if our findings in *E. coli* could extend to other organisms. To answer this question, we retrieved the proteomes of five unicellular organisms with different degrees of phylogenetic proximity to *E. coli*. We chose *Haemophilus influenzae*, which belongs to the same *Gammaproteobacteria* class as *E. coli*; *Helicobacter pylori*, another Gram-negative bacterium, which belongs to a distinct phylum, *Campylobacterota*; *Staphylococcus aureus*, a Gram-positive bacterium of the phylum *Bacillota*; *Methanocaldococcus jannaschii*, a thermophilic archaeon; and *Saccharamoyces cerevisiae*, a eukaryote. Then, we determined the distribution of SASA/L and average hydropathy for the non-membrane proteins in these organisms, distinguishing between proteins of different COG-categories (or arCOG or KOG, for *M. jannaschii* and *S. cerevisiae*, respectively). Across these five organisms, we observed the same pattern as for *E. coli*, with COG-E proteins being significantly more compact (i.e., low SASA/L) and overall more hydrophobic (i.e., higher average hydropathy) than the remaining proteins (Fig. 5K). Thus, similar physical properties characterize proteins involved in amino acid metabolism across domains of life. These findings therefore suggest that the tendency for agglomeration that we identified for *E. coli* COG-E proteins may be evolutionarily conserved.

## Discussion

In this work, we show that the diffusion coefficient of a 40 nm fluorescent particle changes by more than 3-fold in the cytoplasm of *E. coli* across exponential growth on different nutrients. We found that the diffusion coefficient shows an anticorrelation with the abundance of proteins involved in amino acid metabolism (COG category “E”), robust to various perturbations. The observed increase of cytoplasmic viscoelasticity with increased COG-E protein abundance suggests the formation of protein assemblies that obstruct the mobility of the fluorescent particle. Our in vivo and in vitro observations indicate that COG-E proteins can form supramolecular assemblies, namely amorphous agglomerates, likely enabled by their compactness and hydrophobic characteristics (Fig. 6).

**Figure 6.**
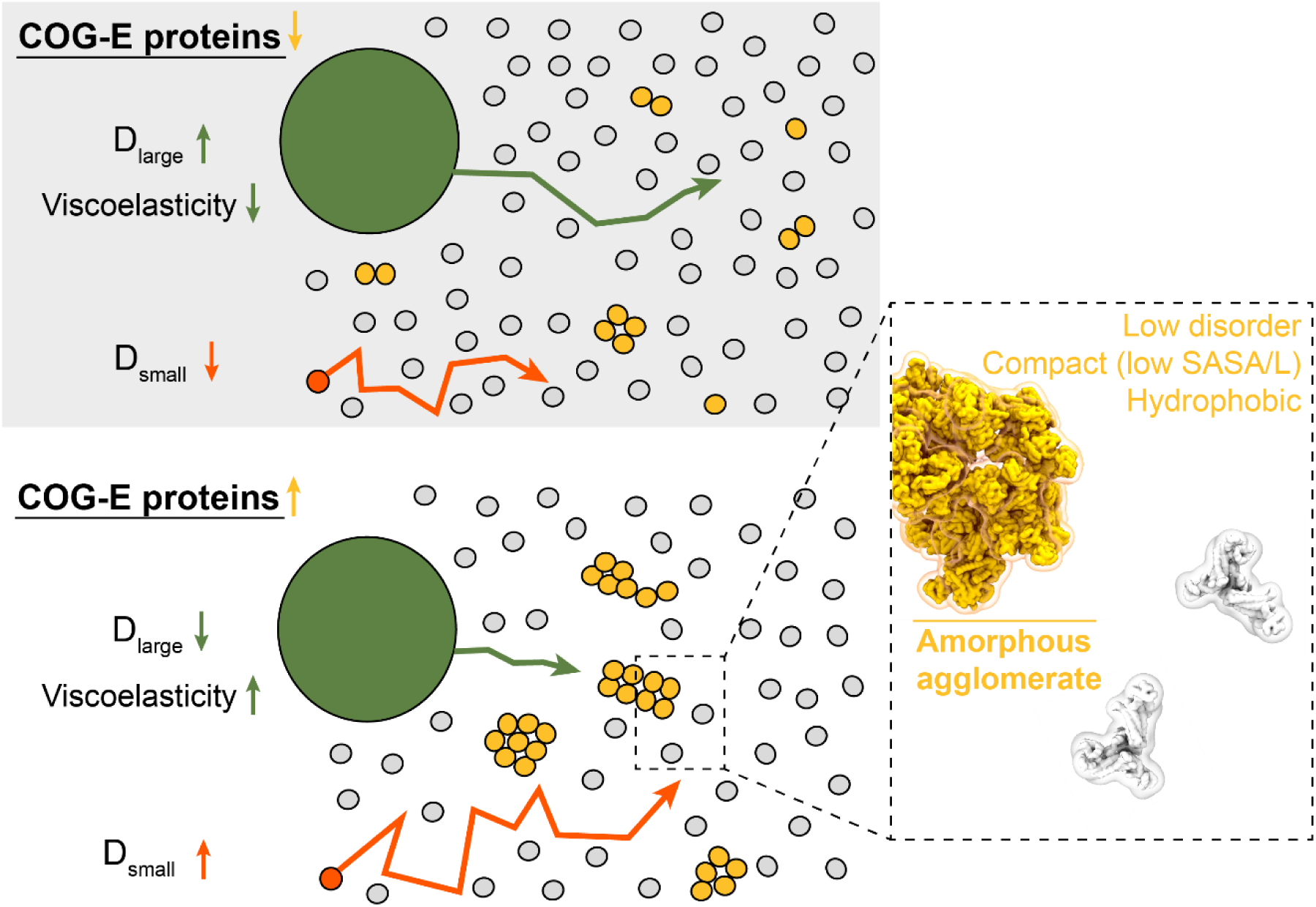
Proposed model for the condition-dependent effect of COG-E protein abundance on diffusion and cytoplasmic organization. Schematic showing reduced protein agglomeration in the cytoplasm when *E. coli* is grown in a condition of low COG-E protein abundance (yellow) relative to other proteins (grey; top left). The second scenario represents a condition in which COG-E proteins are more abundant, and hence agglomeration is more prevalent (bottom left). The presence of protein agglomerates hinders the diffusion of large particles (green) and renders the cytoplasm more viscoelastic, while the reduction of excluded volume enables the faster diffusion of a smaller probes, like mEos3.2 (dark orange). We hypothesize that the formation of amorphous agglomerates is driven by the physical properties of COG-E proteins, namely their low intrinsic disorder, high compactness and hydrophobicity, as shown by the close-up on the right.

The evidence we present led us to conclude that the supramolecular assemblies formed by COG-E proteins are agglomerates. Yet, other types of assembles exist, such as LLPS and aggregates^78^. By bringing together several observations, we can reinstate our conclusions that these assemblies are agglomerates. Specifically, in our EM analyses of purified proteins, we observed that diluted samples had smaller protein assemblies, consistent with a type of structure that forms reversibly in a concentration-dependent manner. This reversibility excludes the possibility of aggregates of misfolded proteins, which are stable in aqueous solution and whose dissolution requires the presence of chaperones, detergents or chaotropic compounds^81,82^. We also observed that the proteins exhibiting this reversible assembly-formation, i.e., COG-E, were not intrinsically disordered, but instead had low disorder scores, unlike many proteins that undergo LLPS^83^. Interestingly, a lack of disordered regions is not only consistent with, but may even explain, the tendency of these proteins to form assemblies^84^. Furthermore, we also observed that most assemblies formed in vitro maintained irregular shapes, which allowed us to confidently conclude these are not liquid-like assemblies but most likely amorphous “agglomerates” of folded proteins^78^.

We suspect that the formation of these agglomerates could be facilitated by the temporal and spatial organization of protein translation in the crowded, far-from-equilibrium environment of the cytoplasm. Indeed, proteins are locally produced: For a protein to be maintained at a high steady-state concentration (e.g., 10^3^ protein copies/cell for a highly abundant COG-E protein), it must be synthesized at a comparably high rate (10^3^ protein copies/h per cell, for a division time of 1h). Since most mRNA transcripts, especially those encoding proteins involved in amino acid synthesis, have half-lives of less than 5 min^85^, and are present in low copy numbers at any given time (i.e., 10^0^ mRNA copies/cell^86^), this implies that translation of such protein occurs at discrete spatial locations and in temporal bursts (10^2^ protein/mRNA). Importantly, for proteins to cluster at the site of translation, successively translated proteins must physically interact before either of them has the chance to diffuse away. Indeed, multiple ribosomes can simultaneously translate the same mRNA transcript (polysomes)^87^, a mode of translation which has been shown to facilitate co-translational assembly of (primarily homo-) oligomers^88–91^. It is tempting to think that translation by polysomes could similarly enable the interactions that lead to agglomerate formation. Moving beyond agglomerates of alike proteins, it is conceivable that such assemblies could include proteins encoded at different gene loci, as mRNAs are known to remain close to the chromosomal positions from which they are transcribed^92^. Overall, mRNAs would be “nucleating agents” for the formation of protein agglomerates, which is consistent with the discrete number of protein clusters in cells which we found with super-resolution microscopy.

Earlier studies have reported that starvation and other forms of metabolic inhibition severely hamper the diffusion of large exogenous probes^25–27^, while leaving smaller probes unaffected^16,25^. While some explained the restricted mobility in such conditions with an increase in cell density^26^, others explained it with the “gelation” of the cytoplasm that results from enhanced protein-protein interactions as the intracellular pH drops^27,93,94^. Indeed, lower pH leads to the neutralization of negatively charged surface residues, thereby reducing the repulsive electrostatic forces between proteins and facilitating agglomeration^78,95,96^. Thus, the action of metabolic inhibitors via a drop in pH could enhance protein agglomeration, beyond the levels found in exponential growth conditions.

A key question that arises from our findings is why COG-E proteins form such agglomerates, i.e., if these agglomerates have a functional role. First, just like random mutations can cause proteins to become entrenched in oligomeric states without loss or gain of function compared to the monomer^97^, it could be that agglomerates exist for “no reason”. On the other hand, COG-E proteins are among the most abundant proteins^54^, and are involved in the synthesis of amino acids, themselves costly cellular building blocks^98^. As such, we suspect that agglomeration could lead to an increase in enzymatic activity, as seen in artificially induced enzyme clustering^99^, thereby providing an evolutionary advantage. Going a step further, if such assemblies do not solely consist of one type of protein, but include enzymes catalyzing sequential steps of the same metabolic pathways, generally encoded in operons, then they could enable increased local substrate concentrations and enhanced reaction kinetics, while also minimizing the release of toxic or labile metabolic intermediates to the cytoplasm, as previously suggested in the context of metabolons^100,101^. Future research is needed to investigate (i) if such heterogeneous clusters form in vivo, (ii) if the enzymes in them experience enhanced activity, and (iii) if the clusters do endow cells with enhanced evolutionary fitness over strains that are incapable of forming them. For now, it is noteworthy that a recent preprint shows that protein-protein interactions are stronger among enzymes of metabolic pathways with labile intermediates, and that such interactions are mediated by residues other than those involved in homo-oligomerization, in a manner consistent with the agglomeration we posit happens among COG-E proteins^102^.

This work has shown that diffusion at the nanometer-scale in the cytoplasm of *E. coli* is a condition-dependent phenomenon, driven primarily by the abundance of COG-E proteins, which are prone to forming agglomerates. Our finding of COG-E protein agglomeration in vivo could mean that metabolic engineering approaches to amino acid biosynthesis may be more successful if the spatial organization of enzymes is considered. More importantly, these results show that the cytoplasm has a previously underappreciated level of spatial organization, establishing a connection between its rheology and the metabolic conditions, which seems to span beyond *E. coli,* as the conserveness of the physical properties of COG-E proteins suggest.

## Materials and Methods

### Strains and probes

The *E. coli* strain BW25113 (*lacI^q^ rrnBT14 ΔlacZWJ16 hsdR514 ΔaraBADAH33 ΔrhaBADLD78*) and corresponding knockout mutants Δ*ptsG* and Δ*argR* from the KEIO collection^103^ were used for most of the experiments described in this work. The strain LY177 (Δ*recA-Tc* ydeO::*I-Sce1^CS^ ilvA::I-Sce1^CS^*), derived from MG1655^104^, was used in experiments for targeted depletion of macromolecules. This strain was kindly provided by Dr. Christian Lesterlin (CNRS-INSERM; Lyon, France).

For single particle tracking, strains were first transformed with a plasmid encoding PfVS, an encapsulin fusion protein that self-assembles into a particle of approximately 40 nm in diameter^29^, under the regulation of an anhydrotetracycline-inducible promoter, *Ptet*. The plasmid was constructed by Gibson Assembly^105^, after amplification of the *PfVS* gene from pRS306-URA3-PHIS3- PfV-GS-Sapphire^29^ by high-fidelity PCR with primers designed with 5’ overhangs complementary to the plasmid backbone (5’-gaaaagaattcaaaagatctATGCTCTCAATAAATCCAAC-3’ and 5’- cctggagatccttactcgagTTATTTGTACAATTCATCAATACC-3’). Leaky protein expression was sufficient for visualization of the particles.

For intracellular pH measurements, cells were transformed with pNTR-SD-pHluorin, a plasmid encoding the pH-sensitive fluorescent protein pHluorin^48^ under the control of an IPTG-inducible promoter, *Ptac*. This plasmid was also constructed by Gibson Assembly, where the pNTR-SD backbone was amplified with primers (5’-ggatgaactatacaaataaTGAGGCCGGGAATTCAGCTATAG-3’ and 5’-TGGCCACCTCCTTAGGATC-3’) that were partially complementary to the primers used to amplify the *pHluorin* insert (5’-ctaaggaggtggccaATGAGTAAAGGAGAAGAAC-3’ and 5’- TTATTTGTATAGTTCATCCATGC-3’).

For single-molecule displacement mapping, we used BW25113 cells carrying either mEos3.2 or ilvC- mEos3.2 in pBAD vectors, used in a previous study^17^.

For super-resolution imaging of proteins in vivo, we constructed, via chromosome integration, strains in which the proteins of interest were tagged on the C-terminal with mEos3.2 and expressed under their corresponding native promoters. Chromosomal integration was performed following the λ-red protocol^103,106^. Briefly, we amplified DNA fragments encoding a GGTGGS linker sequence followed by mEos3.2 (sequence obtained from Śmigiel *et al*^17^), and by a kanamycin-resistance gene, with its own promoter and flanked by FRT regions (obtained from pKD13^106^). These amplified linear

DNA fragments contained 30 bp overhangs for the downstream region of each gene of interest in the chromosome of *E. coli* BW25113 (i.e., *argD* and *ilvI*). The fragments were electroporated into BW25113 cells expressing a helper λ-red recombinase. The kanamycin cassettes were not removed. The constructs were confirmed by sequencing.

### Preculturing and growth conditions

Cells from glycerol stocks (-70 °C) were streaked on LB-agar plates. To ensure that cells were fully adapted to the different carbon sources under analysis, an extended preculturing procedure was used. For each experiment, a colony, taken from a plate, was inoculated into a loosely-capped culture tube with 2 mL of M9 medium (42.2 mM Na2HPO4, 22 mM KH2PO4, 8.6 mM NaCl, 11.3 mM (NH4)2SO4, 1 mM MgSO4, 6.3 µM ZnSO4, 7 µM CuCl2, 7.1 µM MnSO4, 7.6 µM CoCl2, 100 µM CaCl2, 60 µM FeCl3, 2.8 µM thiamine hydrochloride) supplemented with the appropriate carbon source concentration (see below) and incubated overnight at 37 °C, with orbital shaking at 300 rpm. The following morning, the cell concentration was determined by flow cytometry (BD Accuri C6) and the cells diluted into a 100 mL flask containing 10 mL of preheated medium. The dilution was adjusted based on the known growth rate, so that cells would reach a concentration of ∼1.5 – 3.0 x 10^8^ cells/mL (or an OD600 nm≈0.3) after an 8 – 9 hour incubation. A similar dilution was performed after this period, for an additional step of overnight growth. Cells were sampled for the experiments when the target concentration was reached.

The M9 medium was supplemented with one of different carbon sources, at the following concentrations, as used by Schmidt *et al*^54^: 2.2 g/L glycerol; 2.3 g/L galactose; 5 g/L (D)-glucose; 5 g/L mannose; 5 g/L fructose; 3.3 g/L sodium pyruvate; 3.5 g/L sodium acetate; 2.8 g/L sodium fumarate; 5.7 g/L disodium succinate. Where mentioned, the medium was further supplemented with casamino acids (Formedium) at 2 g/L or L-arginine at 100 µg/mL.

### Microscopy imaging

Upon reaching the desired cell concentration, samples were collected in 0.22 µm centrifuge tube filters (Costar), centrifuged at 11,000 g for 30 s and concentrated 5 to 10-fold by resuspension in a fraction of the flow-through. Two microliters of the concentrated cell suspension were spotted on a glass coverslip (#1.5 thickness, 0.16 – 0.19 mm; Epredia) and were quickly covered with an agarose pad preheated to 37 °C. These pads had the same medium composition as the one used in the preculturing steps and additionally 15 g/L agarose. The samples were rapidly transferred to an incubator housing the microscope likewise preheated to 37 °C.

All single particle tracking and intracellular pH measurements were performed using an Eclipse TiE inverted microscope (Nikon Instruments) and a 100x Plan Apo Lambda NA 1.45 objective (Nikon Instruments), with additional 1.5x optical magnification. The system was coupled to an iXon Ultra 897 EMCCD camera (Andor) and an LED light source (Aura Light II, Lumencor). Two filter sets were used, both containing a 495 nm dichroic mirror and a 525/50 nm emission filter, and either a 390/40 nm or 470/40 nm excitation filter (AHF Analysentechnik). The first set was used for all measurements done with incoming light at 395 nm, and the second set for all measurements done with incoming light at 485 nm, as detailed below.

For single particle tracking, cells were first excited with 395 nm (100% intensity) for 2 s to increase the signal-to-noise ratio, as we serendipitously found this step to have a photoactivating effect on the Sapphire fluorophore. Excitation was then switched to 485 nm (100% intensity), and 100 frame-long movies were acquired using an exposure time of 19.4 ms per frame, camera readout at 17 MHz and EM gain of 100. Cells were also imaged with the brightfield channel for segmentation purposes.

For intracellular pH measurements, snapshots of cells were taken under excitation at 395 and 485 nm. In both channels, the intensity was set to 10%, the exposure time was set to 100 ms and no gain was used.

For single-molecule displacement mapping (SMdM) and photoactivated localization microscopy (PALM), a customized inverted wide-field IX-81 microscope (Olympus) was used, equipped with a 100x objective NA 1.49 (Olympus) and an ET 605/70 M bypass filter (Chroma), coupled to EM-CCD camera (C9100-13, Hamamatsu). The imaging conditions were the same as described earlier^17^. In brief, at the beginning of every other movie frame, a low-intensity laser pulse at 405 nm (OBIS 405 LX, 50 mW maximum power) was used to photoconvert a few mEos3.2 proteins from green to red-fluorescent state. Afterwards, two high-intensity laser pulses at 561 nm (OBIS LS 561-150) were applied to visualize the photoconverted proteins. These pulses were applied on consecutive movie frames with time separation of Δt=1.5 ms such that the displacement of the proteins within this short time period could be captured despite the lower time resolution of the camera (frame rate approximately 18 ms). For each measurement, movies of either 10,000 (PALM) or 100,000 (SMdM) frames were acquired.

### Cell segmentation

Cell segmentation was done using brightfield snapshots. For this, we first applied a rolling-ball background subtraction in Fiji^107^, with *radius*=*20 pix* and the “light background” option enabled. We then used the *MicrobeJ* plug-in^108^ with the following criteria: *area [µm^2^] = [0.5 - 5]; length [µm] = [1.5-max]; width [µm]=[0.3 - 1.6]; angularity amplitude=[0 - 0.6]*; remaining parameters were set to *[0 - max]*. Manual adjustments were performed to eliminate incorrectly segmented cells. Furthermore, cells were labeled according to the presence/absence of visible constriction rings.

### Single particle tracking

Single particle tracking and determination of the mean squared displacement, *MSD*, of each particle was done by sequentially using the functions *batch()*, *link()*, *filter_stubs()* and *imsd()* of the *trackpy* package in Python^39^. Particle localization was done using the function *batch()*, with parameters *diameter=9*, *minmass=0*, *preprocess=True*, *noise_size=1*. Trajectories were obtained with the function *link()*, with parameters *search_range=5*, *memory=1*, *adaptive_step=0.8*. The function *filter_stubs()* was used to select trajectories with a minimum length of *threshold=33* frames. The *MSD* of each particle was obtained with *imsd()*, with parameters *mpp=0.1067*, *fps=51.61*, *max_lagtime=33*. An automatic filtering step was then applied to remove spurious trajectories based on the properties reported by the tracking routine, namely the shape (mean *eccentricity* < 0.2), size (mean *size* < 2.1) and precision of localization (median *ep* < 0.31) of the detected particles.

Cells and corresponding particle trajectories were rotated to a common orientation (long cell-axis parallel to the x-axis, and short cell-axis parallel to the y-axis) using the *ColiCoords* package^109^. Thereafter, one-dimensional mean squared displacements, *MSDx* and *MSDy*, were obtained using either the longitudinal or radial coordinates of the particle trajectories, respectively.

Diffusion coefficients were obtained for every tracked particle by fitting the *MSD* to one of the following equations:

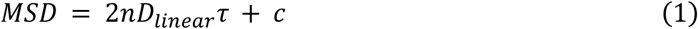

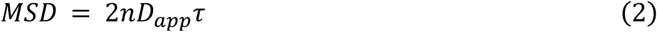

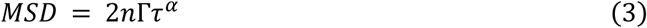

where *Dlinear, Dapp and* Γ are the linear, apparent and generalized diffusion coefficients, respectively; *n* is the number of dimensions (*n* = 1, in the case of one-dimensional diffusion); *α* is the anomalous diffusion exponent; and *c* is an offset, generalized used to account for errors in the localization^110^. Except where noted, the values of diffusion coefficient reported in this work refer to the fitting of *MSDx* to Eq. 1, i.e., to the values of *Dx, linear*.

### Post processing of particle tracks

The brightness of each particle was defined as the median of the brightness ("mass" values obtained from *trackpy*) in the first five frames in which that particle was found. To remove potential aggregates or clusters of multiple particles that were detected as one, we removed "particles" with values of brightness above certain cutoff values. We used two approaches, differing in on how these cutoff values were defined: (i) with the “fixed filter” approach, we removed all particles having log10(brightness) higher than 3.5, a value which we chose after visual inspection of the distributions of particle brightnesses across many conditions; (ii) with the "adaptive filter" approach, we applied a condition-dependent cutoff value, defined separately for each growth condition as median{log10(brightness)} + 0.25. Throughout the main text, we used the "adaptive filter" approach. In Fig. S1C, the two methods are compared.

### Particle trajectory simulations

To simulate particles moving with Brownian motion in cell-shaped compartment, we used the Python API of *Smoldyn*^41^. First, the spherocylindrical compartments were obtained by combining a cylinder and two hemispheres of radius 0.4 µm. To emulate cells with different lengths, simulations were performed where the length of the cylinder was varied between 0.3 and 3.6 µm. Then, 500 particles were added to the compartment as a single “species”, with a defined diffusion coefficient. For each simulation, this diffusion coefficient of the species was fixed to a value between 0.02 and 0.2 µm^2^/s. The interaction between the compartment surface and the particles was set to be fully reflective, by setting *setAction*() with parameters “both” and “reflect”.

### Analysis of particle trajectories with ExTrack

Particle trajectories were analyzed with *ExTrack*, a tool that fits a hidden Markov model to the observed particle data^66^. We used this package to estimate the diffusion coefficient and fraction of each diffusive state, as well as the transition rates between states, for each growth condition. The localization error was likewise estimated with this package.

We used a three state model to analyze the data, as the likelihood of this model, *L3*, was significantly higher than the likelihood of a two-state model, *L2* (*L2*/*L3* < 1 x 10^-185^). When running 2-state or 4-state models, we found qualitatively similar results to the 3-state model, namely, an immobile/slow diffusion state and faster diffusive states whose mean diffusion length was anticorrelated with COG- E density (*r^2^* >70% for all models from 2 to 4 states; comparable to Fig. 3I).

The state with the lowest diffusion coefficient, i.e. state “0”, was designated as the “slow” state. The remaining states were designated as “fast” states. For every growth condition *j*, we obtained a representative diffusion coefficient for these “fast” states by averaging the diffusion coefficient of each state, *Dij,* weighted by its fraction, *fij*:

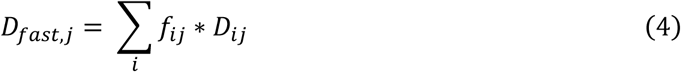

### Single Molecule Displacement Mapping (SMdM)

Blinking events of mEos3.2 fluorophores in cells were detected using a previously developed package for the analysis of STORM data^111^. To establish the limits of each analyzed cell, the 2D-coordinates (x, y) of the detected fluorescent peaks were clustered using the Voronoi tessellation method with a minimum cluster size of 2000 and a minimum cluster density of 0.35 times the median density. This density was defined as the inverse of the area of a polygon surrounding the cloud of points. Then, protein displacements were obtained as the Euclidean distance between the fluorescent peaks in consecutive movie frames, with each displacement starting on an odd-numbered frame and ending on an even-numbered frame. All possible displacements within a search radius of 600 nm were measured.

The distribution of fluorescent protein displacements was fitted with an adjusted probability density function of a 2-dimensional random-walk diffusion model, corrected with a linear term to account for background noise, and extended to allow for multiple diffusive states^17,112^:

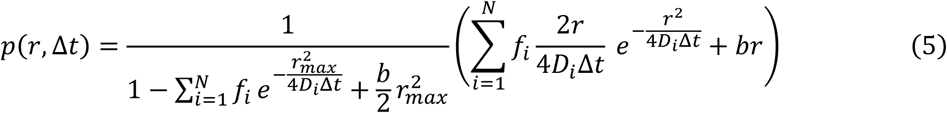

where 𝑟 is the displacement of a fluorescent protein between consecutive 561 nm readout laser pulses; 𝑟_𝑚𝑎𝑥_ is the maximum search radius (600 nm); Δ𝑡 is the time between these laser pulses (1.5 ms); 𝑏 is a background correction factor; 𝑁 is the number of diffusive states (assumed to be 1); 𝐷_𝑖_ is the diffusion coefficient of state 𝑖; 𝑓_𝑖_ is the fraction of particle of displacements of state 𝑖, such that 𝑓_𝑖_ = 1. Here, 𝑏, 𝐷_𝑖_ and 𝑓_𝑖_ are parameters to be estimated in the maximum likelihood estimation fitting of Eq. 5 to the distribution of protein displacements.

### Photoactivated light microscopy (PALM)

For the reconstruction of super-resolution images, we used the function *plot*_*storm*() from *ColiCoords* with settings *method*=*"hist"* and *upscale*=*1*. This was done either for individual cells, or for multiple cells. In the latter case, we used the function *align*_*data*_*element*() and mapped the reconstructed images onto a synthetic "cell", *SynthCell*(), with *length*=*30* and *width*=*8*.

For clustering of blinking events, we used *DBSCAN* algorithm from *scikit-learn*, with parameters *eps*=*0.3*, *min_samples*=*10*.

### Intracellular pH measurements

The camera baseline value was subtracted from the raw microscopy images of cells expressing the pHluorin sensor. For each fluorescence channel, we then obtained the mean fluorescence intensity of each cell using the function *measure.regionprops_table()* from the *scikit-image* package in Python^113^. The readout of the pH sensor pHluorin was defined as the ratio between the mean intensity obtained under excitation at 395 nm and the mean intensity under excitation at 485 nm. BW25113 cells without pHluorin were analyzed in the same way to enable for the correction for autofluorescence.

### Determination of features of E. coli proteins

We used the published proteomics data set (Schmidt *et al*^54^; Table S6 therein) as reference in this work. This data set includes 2359 proteins, of which 2357 were present in at least one of the ten growth conditions that we used for the single particle tracking experiments (i.e., minimal medium with acetate, fumarate, fructose, galactose, glucose, glycerol, glycerol with casamino acids, mannose, pyruvate, succinate). We used the UniProt accession codes and COG categories as provided in the data set. In the 200 cases where multiple COG categories had been assigned to a single protein, only the first category was considered (e.g., a protein assigned to category “EG” in Schmidt *et al*’s data set was re-assigned to category “E”).

The provided UniProt accession codes were used to automatically retrieve data for each protein from the UniProt database^114^ (accessed on 13/07/2023) using a custom-made script. The *location* was obtained from the sub-fields “comments > SUBCELLULAR LOCATION” and “uniProtKBCrossReferences > database > GO”. Each protein was then considered *cytoplasmic* if only the terms “cytosol” or “cytoplasm” were found; or as *external* if at least one of the terms “membrane”, “surface”, “periplasm”, “fimbrium”, “flagellum” or “secreted” was found.

To determine structure-based protein features, we used the 3D structures of *E. coli* proteins retrieved from the AlphaFold Structure Database^115,116^ (accessed on 4/10/2022). Structures were available for 2341 proteins. The *MDTraj* package^77^ was used to obtain descriptors of the overall shape of these structures, namely asphericity, relative shape anisotropy and radius of gyration. We also used this package’s implementation of the Shrake-Rupley algorithm^117^, with default parameters, to obtain the solvent-accessible surface area of each amino acid, 𝑆𝐴𝑆𝐴_𝑖_. The values of 𝑆𝐴𝑆𝐴_𝑖_ were normalized by the maximum solvent accessibility of each amino acid, 𝐴𝑆𝐴_𝑖_^118^, to give the relative solvent-accessible area of that amino acid, 𝑟𝑆𝐴_𝑖_:

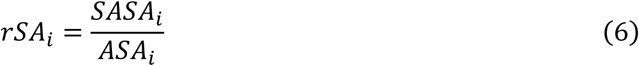

Residues with 𝑟𝑆𝐴_𝑖_ > 0.25 were classified as “exposed”.

We determined both the *total* solvent accessible surface area and the *hydrophobic* solvent accessible area, by summing 𝑆𝐴𝑆𝐴_𝑖_ either over all 𝑖 residues or over 𝑖 𝜖 {𝐴, 𝐶, 𝐹, 𝐼, 𝐿, 𝑀, 𝑉, 𝑊, 𝑌}, respectively. The ratio between these two areas yielded the relative exposed area of hydrophobic residues. To determine the areas of the three largest hydrophobic patches, we used *MolPatch*^76^.

For sequence-based protein features, we used (i) the functions *secondary_structure_fraction*(), *aromaticity*(), *isoelectric_point*() and *get_amino_acids_percent*() of the *Biopython* package^72^; (ii) the function *GetCTD*() of the *PyBioMed* package^73^; (iii) the *NetSurfP3.0* web-server^75^; (iv) and *IUPred2A*^74^. Since the last two tools provide disorder score values per amino acid, we further determined an average value across the entire sequence, as well as the length and number of stretches of more than 10 adjacent amino acids with an index > 0.5. The “disorder score” mentioned in the main text corresponds to the sequence-average of values obtained with *NetSurfP3.0* (c.f. “_nsp3_disorder” in Table S1).

As a measure of the overall protein hydrophobicity, we determined the average hydropathy score, according to the Kyte-Doolittle scale^119^, over all residues. As a measure of surface hydrophobicity, we averaged the hydropathy score over solvent-exposed residues (i.e., residues with *rSA* > 0.25).

### Supervised machine learning models for binary classification

Reasoning that the diffusion of the cytoplasmic fluorescent particle is affected by highly abundant proteins in that compartment, we excluded proteins assigned to an *external* location (773 proteins in total) and those with an average abundance lower than 100 copies/cell (1049 proteins in total). This led to a reduction of the original data set to 899 proteins.

We then defined labels for each protein according to the observed correlation between their abundance and the experimentally measured diffusion coefficients. Labels “non-correlating” and “anticorrelating” were assigned to proteins for which these correlations were described by |*r*| < 0.3 and *r* < -0.5, respectively. Next, all proteins that either lacked a label (i.e., proteins with *r* > 0.3 or with -0.5 < *r* < -0.3), or missed some feature values were excluded, further reducing the data set to 465 proteins (of which: 81 anticorrelating and 384 non-correlating).

To avoid pairwise collinearities among the features, we first determined the Pearson correlation coefficients, *rfeature pair*, for all pairs of features in our protein training set (as defined below). We then excluded the second feature from every pair for which |*rfeature pair*| > 0.8. Then, to identify and remove multicollinearities, we determined the Variance Inflation Factor, *VIF*, for each feature, and iteratively dropped all features for which *VIF* > 5. Overall, and after removal of collinear features, we obtained 76 or 11 sequence- and structure-based features (Tables S1 and S2).

The data was split using *train_test_split*() from the *scikit-learn* package in Python^120^ with *test_size*=*0.2* and stratification according to the class labels, yielding training and test sets of 372 and 93 proteins, respectively. This process was repeated four additional times with different seeds, each generating different train-test splits (referred to as “data folds” in Fig. 5F).

We implemented a two-step machine learning pipeline which first scales the feature values with *RobustScaler*() and then performs the classification with a *RandomForestClassifier*() estimator. Hyperparameters were set to: *n_estimators=100, max_depth=3, criterion=“gini”, max_features=“sqrt”* and *class_weight*=*“balanced”*. During training, the F1-score of the positive class (label “anticorrelating”) was used as the scoring function. To account for the stochasticity in model training, we repeated this training step n=50 times on the same training data set.

Among various models trained on the same dataset, we selected those with the highest value of AUC-ROC score obtained during training. Then, we obtained a non-biased measure of the model performance by determining the AUC-ROC score on the unseen test set (93 proteins). This test set was also used to determine measures of feature importance with *permutation_importance* from *scikit-learn,* with *n_repeats=50* and the scoring function set to the F1-score.

### Analysis of the proteomes of other organisms

We obtained the proteomes of five organisms, *Haemophilus influenzae* (“*Hi*”, UP000000579)*, Helicobacter pylori* (“*Hp*”, UP000000429)*, Staphylococcus aureus* (“*Sa*”, UP000008816)*, Methanocaldococcus jannaschii* (“*Mj*”, UP000000805) and *Saccharomyces cerevisiae* (“*Sc*”, UP000002311) from UniProt (accessed on 31/03/2025), as well the corresponding 3D structure predictions from the AlphaFold Structure Database (accessed on 31/03/2025). We determined the solvent accessible surface area normalized by sequence length, SASA/L, and the average hydropathy for each protein as described above. We further excluded *external* proteins from our analysis as described above.

To assign COG categories to each protein, we first used the “Retrieve/ID mapping” tool of UniProt, mapping the proteins’ primary accession codes to the IDs used in the eggNOG database. As a result, most proteins were assigned a COG (for *Hi, Hp* and *Sa*), arCOG (for *Mj*), or KOG ID (for *Sc*). Some proteins were instead assigned to categories that are specific to this database (“ENOG…”). The fractions of proteins with the latter type of categories were 3.3%, 3.6%, 10.7%, 0.1% and 27.4% for *Hi*, *Hp*, *Sa*, *Mj* and *Sc*, respectively. The functional categories of all COG/arCOG/KOG IDs, were downloaded from the COG database (NCBI^121^; accessed on 01/04/2025). The remaining proteins were not assigned to any functional category.

### RNAseq

For reverse transcription, 2 µg of total RNA were mixed with 1 µL 20 µM random 9N primers, 1 µL TGIRT-III RT (InGex), 4 µL 5X RT Buffer (250 mM Tris pH 8.3, 375 mM KCl, 15 mM MgCl2), 1 µL dithiothreitol 0.1 M and1 µL dNTPs (10 mM each), in a total reaction volume of 20 µL and incubated at 25 °C for 10 min, 57 °C for 1 h.

To convert the RNA-DNA hybrid to dsDNA, the TGIRT-III enzyme was first degraded by addition of 1 µg Proteinase K, incubating at 37 °C for 20 min. Then Proteinase K was inactivated by addition of 0.5 µL of Sigma Aldrich Protease Inhibitor Cocktail. The reaction was used as input for the NEBNext Second Strand Module, and second strand synthesis was performed at 16 °C for 1 h, as per manufacturer instructions.

Finally, dsDNA was then tagmented using the Illumina Nextera XT kit, as per manufacturer instructions.

### Proteomics

Upon reaching a target cell concentration as used for single particle tracking experiments, cells (in biological triplicates) were collected by centrifugation and washed twice with PBS at 4 °C. Centrifugation was done at 20,000 g, at 4 °C, for 1 min. The cell pellets were then flash-frozen in liquid nitrogen and stored at -70 °C until further processing.

To perform cell lysis, the pellets were resuspended in lysis buffer (2% sodium deoxycholate in 100 mM ammonium bicarbonate) and 1 µL of reducing agent solution (0.2 M tris(2-carboxyethyl) phosphine, TCEP, in 100 mM ammonium bicarbonate) was added per 40 µL of cell suspension. After vortexing for 10 s, the samples were sonicated with a VialTweeter (Hielscher; two cycles of 10 s, at maximum amplitude), and heated to 95 °C in a thermomixer for 10 min at 500 rpm. Small aliquots of each lysate sample were taken for protein quantification using the Pierce™ Microplate BCA Protein Assay Kit (ThermoFisher Scientific), following the manufacturer’s instructions.

After quantification, samples containing 50 µg of protein were denatured with 1 M urea and reduced with 10 mM TCEP for 1 h at 37 °C, and then alkylated with 15 mM iodoacetamide at room temperature, in the dark, for 45 min. Subsequently, these samples were digested by 1:50 (w/w) sequencing grade modified trypsin at 37 °C overnight with 15 s shaking at 800 rpm for every 5 min. After digestion, trifluoroacetic acid, TFA, was added to a final concentration of 1% (v/v) and the precipitated detergent was separated by centrifugation. The digested peptides in the supernatant were then cleaned with C18 tips (Pierce) according to the manufacturer’s instructions. The cleaned peptides were dried by centrifugation under vacuum and reconstituted in 20 µL of a solution of 2% acetonitrile and 0.1% formic acid, FA. Peptide content was measured by the absorbance of 280 nm.

The reconstituted peptides (1.2 µg) were loaded into a C18 Trap cartridge (Thermo Acclaim PepMap C18 Reversed Phase Trap Cartridge, 5 µ, 0.3 mm I.D. x 5 mm L.) and separated by reverse phase chromatography using a 75 µm I.D. x 15 cm length nano-LC column (Thermo Acclaim PepMap RSLC C18 2 µ). Peptides were eluted with a linear gradient starting from 97% solvent A (0.1% FA) and 3% solvent B (80% acetonitrile and 0.1% FA) to 35% solvent B, over 60 min, at a flow rate of 300 nL/min by Ultimate 3000 RSLC chromatography system (ThermoFisher Scientific). The eluted peptides were ionized by online nano-electrospray and measured by Exploris 480 Mass analyzer (ThermoFisher Scientific). Data dependent acquisition mode was used to obtain high resolution master scans from 385 m/z to 1540 m/z at a resolution of 120,000 at m/z 200. Ions at +2 to +6 charge states were selected and the ions with unassigned charge state were excluded. The 20 most intense selected ions were then fragmented by HCD at 30% normalized collision energy, and the fragment ions were measured by data-dependent, dd, MS2 scans at the resolution of 15000. Dynamic exclusion of the same precursor ions was set to 20 s and ion accumulation time was set to auto for both master scan and ddMS2 scan.

LC-MS/MS raw data was analyzed by MaxQuant v.2.3.1.0 and searched against *E. coli* pan-proteome database downloaded from the UniProt database (Proteome ID: UP000000625, 123487 entries; downloaded on 14/12/2022) using the integrated Andromeda search engine^122,123^. The default instrument parameters for Orbitraps in MaxQuant were used. The following search parameters were used: digestion was defined as enzyme-specific (trypsin), with a maximum of three missed cleavages; carbamidomethylation was set as a fixed modification on cysteines; oxidation of methionines, deamidation of asparagines and glutamines, and acetylation of peptide N-termini were all set as variable modifications; the identification criteria was at least one peptide match; both peptide and protein false discovery rate, FDR, were limited at 1%. The MaxLFQ algorithm was applied for label-free quantification, LFQ^124^ and the time window of match between runs was set at 0.7 min. The quantification values were reported as iBAQ values.

To convert the iBAQ values into estimates of the absolute concentration of each protein, 𝑖, in each condition, 𝑗, we first determined the fold change, 𝑓_𝑖, 𝑗_, of the iBAQ metrics with respect to a reference condition (wild-type strain grown in minimal medium with glucose, “WT-Glc”):

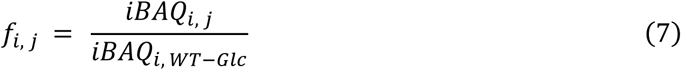

With absolute protein abundances having been previously reported for this reference condition^54^, 𝑐𝑖, 𝑊𝑇−𝐺𝑙𝑐 (𝑆𝑐ℎ𝑚𝑖𝑑𝑡2016), the fold change was multiplied by these reported concentration values to arrive at the concentration in each of the new conditions, 𝑐_𝑖, 𝑗_:

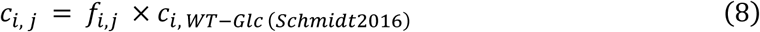

### Protein expression

Genes encoding *argD, argE, argG, aroF, ilvH, ilvI, ilvN, serA, tauD, trpA, trpB* and *tyrA*, codon-optimized for *E. coli* with BsaI sites positioned at the 5′ and 3′ prime ends (Twist Bioscience), were inserted into either pBADHis or pBADHis-SUMO vectors using the Golden Gate methodology^125^. For transformation, 3 μL of the plasmid were added to 50 μL of NEB10β RbCl competent cells and incubated on ice for 30 min. Cells were then subjected to heat shock at 42 °C for 40 s, followed by another incubation on ice for 2 min. Subsequently, 250 μL of prewarmed LB medium were added, and cells were incubated for 75 min at 37 °C. A 50 μL aliquot of cells was plated on LB-agar supplemented with 50 μg/mL ampicillin and incubated overnight at 37 °C. Plasmid isolation was performed, and the success of cloning was confirmed through sequencing.

For protein expression, a 5 mL preculture (LB with 50 μg/mL ampicillin) was incubated overnight at 37 °C, and then used to inoculate 250 mL baffled flasks containing 50 mL of Terrific Broth medium supplemented with 50 μg/mL ampicillin. The flasks were incubated at 37 °C until an OD600 of 0.6−0.8 was achieved. Expression was induced with 0.2 g/L L-arabinose, and cultures were maintained at 24 °C for 18 h before harvesting. Cell harvesting was conducted by centrifugation (3700 rpm, 20 min, 4 °C). Expression was confirmed by SDS-PAGE (BioRad SurePage) and positive samples were selected for further purification.

### Protein purification

Cell pellets were suspended in lysis buffer (50 mM Tris-HCl and 150 mM NaCl, pH 8). Subsequently, 0.10 mM phenylmethylsulfonyl fluoride and 10 μg/ml DNAseI were added to the lysis solution to prevent protein degradation. Cells were disrupted by sonication (5 s “on”, 7 s “off”, 70% amplitude, totaling 5 min) using a Vibra-Cell™ VCX 130 sonicator (Sonics Materials) equipped with a 3 mm stepped microtip, and then centrifuged at 8,000 g for 1 h at 4 °C. The resulting supernatant was applied to a gravity column containing 0.5 mL of Ni Sepharose resin (GE Healthcare), previously equilibrated with lysis buffer, and incubated with the resin for 1 h at 4 °C. After a washing step with five column volumes of wash buffer (50 mM Tris-HCl, 150 mM NaCl and 20 mM imidazole, pH 8), the protein was eluted in two column volumes of elution buffer (50 mM Tris-HCl, 150 mM NaCl, 500 mM imidazole, pH 8). The elution buffer was subsequently exchanged with storage buffer (50 mM Tris, pH 8). If necessary, 5 mM dithiothreitol was added as reducing agent, and 5% (v/v) glycerol was added to increase protein stability. Where applicable, the SUMO-tag was cleaved by SUMO-protease. Protein concentration was determined by measuring the absorbance at 280 nm and the samples stored at 4 °C.

### Mass photometry

Microscope coverslips (24 x 50 mm; Fisher Scientific) were soaked in a solution of 3% mucasol detergent for 30 min. The coverslips were thereafter thoroughly washed by rinsing three times with HPLC-grade ethanol, followed by rinsing three times with MilliQ water. The coverslips were then dried with a stream of nitrogen.

Mass photometry measurements were performed as previously described^126^. First, a 6-well silicon gasket was positioned onto a glass coverslip, which was then inserted in a Two MP mass photometer (Refeyn). Before conducting measurements, the instrument was calibrated using a NativeMark™ Unstained Protein Standard (Invitrogen). Each protein sample was diluted in storage buffer to a final concentration of 10 – 100 nM immediately prior to a measurement session. For each measurement, 8 μL of buffer (pre-filtered through a 0.22 μm syringe) were first added to a well. Then, this volume was finely adjusted using the droplet dilution mode to attain optimal focus. Upon achieving focus, 8 μL of protein sample were added to the same well and gently mixed by pipetting up-and-down. After mixing, data collection was initiated and continued for 1 min. Data acquisition was controlled with the AcquireMP software (Refeyn). Data processing and analysis were conducted with the DiscoverMP software (Refeyn).

### Transmission electron microscopy imaging

For imaging of cell cross sections, *E. coli* cells were concentrated by centrifugation and the pellet was transferred to a 3 mm gold plated type B carrier. Cells were immobilized with high pressure freezing (EM ICE, Leica) and then freeze-substituted in 1% osmium tetroxide and 0.5% uranyl acetate in acetone with 5% water using the quick freeze substitution method^127^. Samples were embedded in Epon and ultra-thin sections were collected on formvar-coated and carbon evaporated copper grids and inspected using a TALOS L120C transmission electron microscope (TEM), at 120 kV.

For imaging of purified protein samples, protein solutions were pipetted onto 400 mesh copper grids coated with a carbon-layer. Excess volume was removed with a paper filter. Uranyl acetate solution at 2% was added to the same grid, and incubated for approximately 1 min. After blotting the stain, the grids were dried and the samples were imaged with either a Talos L120C (ThermoFisher Scientific) or CM120 (Phillips) TEM, at 120 kV.

The sizes of the structures observed in these TEM images were determined by manual measurement in Fiji^107^. For this, we drew straight lines across each structure, along the direction in which edge-to-edge distance appeared to be maximal.

### Computational prediction of homodimer formation

Amino acid sequences were obtained from the UniProt database. Fasta files with two repeated entries of each amino acid sequence were then provided as input to *AlphaFold Multimer* v.2.1.1 or v.2.3.1^80^ for homodimer predictions, using *max_template_date* = 31/12/2021. Each of the five *AlphaFold Multimer* models was initialized with either one or five random seeds, resulting in a total of 5 or 25 predictions per protein, respectively. We used one random seed for 715 predictions and five random seeds for 1156 predictions, respectively. From a total of 1871 protein homodimer predictions (of which 193 COG-E proteins), we then filtered for cytoplasmic proteins, resulting in a dataset of 824 proteins (of which 105 COG-E proteins).

## Resource availability

### Lead contact

Requests for further information and resources should be directed to and will be fulfilled by the lead contact, Matthias Heinemann.

### Materials availability

All plasmids and strains generated in this study are available from the lead contact without restriction.

### Data and code availability

Proteomics data have been deposited at ProteomeXchange Consortium via the PRIDE partner repository with the dataset identifier PXD064526. RNAseq data have been deposited at ArrayExpress with accession number E-MTAB-15217. *AlphaFold* homodimer predictions, original electron microscopy images, single-particle tracking data and figure source data have been deposited at DataverseNL with identifier doi: 10.34894/SQXHBD. Original code is available on GitHub (https://github.com/molecular-systems-biology/Losa-et-al-2025). Any additional information required to reanalyze the data reported in this paper is available from the lead contact upon request.

## Supporting information

Supplemental items

## Acknowledgments

We thank Hein Wijma for helping with the *AlphaFold Multimer* predictions; Silke Bonsing-Vedelaar for constructing the pNTR-SD-pHluorin plasmid; all members of the Molecular Systems Biology group, as well as Alexander Belyy, Arjen van der Schaaf, Daniele Parisi, Dea Gogishvili, Georg Hochberg, Kasia Tych, Marten Chaillet, and Yulia Yancheva for helpful discussions and suggestions; the Center for Information Technology of the University of Groningen for their support and for providing access to the Hábrók high performance computing cluster, as well as Calcul Québec (calculquebec.ca) and the Digital Research Alliance of Canada (alliancecan.ca).

This research was supported by the “BaSyC – Building a Synthetic Cell” Gravitation grant (024.003.019) of the Netherlands Ministry of Education, Culture and Science (OCW) and the Dutch Research Council (NWO) (to M.H.), by the Dutch Research Agenda (NWA) grant (NWA.1292.19.170) financed by the Dutch Research Council (NWO) (to M.H., B.P., S.J.M., M.W.F.), by the Natural Sciences and Engineering Research Council of Canada (NSERC Discovery grant (RGPIN-2021-03208 to S.v.T., RGPIN-2022-05142 to L.E.W.)). S.v.T. and L.E.W. are recipients of salary awards from the Fonds de Recherche du Québec – Santé (FRQS).

## Author contributions

J.L. performed all single particle tracking and epifluorescence microscopy experiments, computational work and data analysis, under the supervision of M.H.; J.L. and D.L. performed SMdM and PALM microscopy, under the supervision of B.P.; F.S. performed the analysis with *ExTrack*, under the supervision of L.E.W. and S.v.T.; S.G.K. performed the protein purification and mass photometry measurements, under the supervision of M.W.F.; J.S. performed computational work under the supervision of S.J.M.; A.S., M.C.A.S. and R.d.B. performed the electron microscopy analyses; F.Y.H. performed the experimental proteomics analysis; D.I. performed the RNAseq analysis; J.v.E. performed computational analysis with *MolPatch*; S.A. supervised the machine learning-based investigations; A.G. advised the investigation of protein properties; J.L. and M.H. conceptualized the study and wrote the manuscript.

## Supplemental information

Document S1. Tables S1-S3.

