## Supplemental items for "Condition-dependent, amorphous protein agglomerates control cytoplasmic rheology"

### Supplemental figures

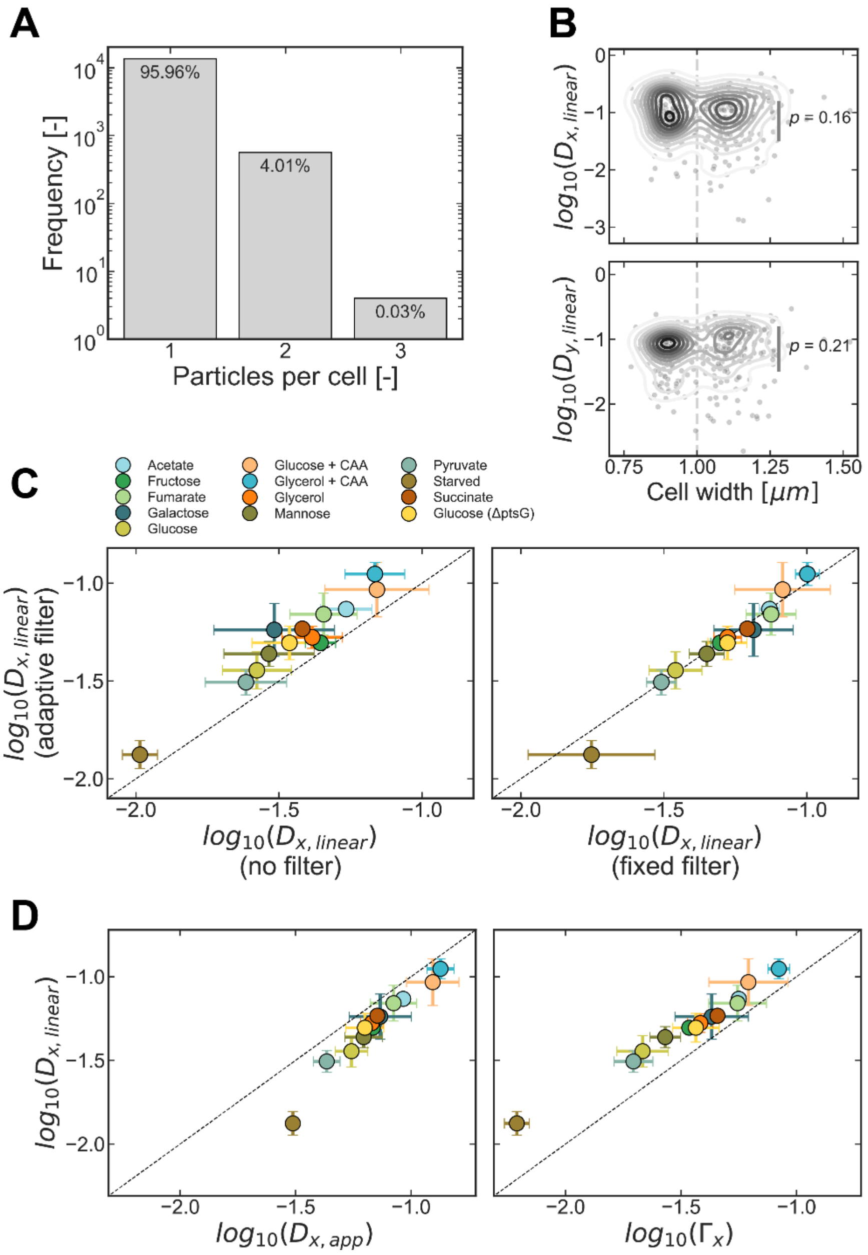


**Figure S1 – Diffusion coefficients of individual particles show condition dependent trends that are robust to variations in cell geometry, post-processing steps and diffusion models used, related to Figure 1.**

(**A**) Bar plot showing the distribution of the number of particles per cell, for all cells and growth conditions shown in Fig. 1B, considered in the analysis. Although uncommon, some cells contained more than 3 particles. In those cases, tracking was impossible due to the proximity of the different particles.

(**B**) Density plots of the diffusion coefficients measured in individual cells grown in medium with galactose, plotted against the width of each cell. This plot shows, for every detected particle, the width of the cell within which that particle was found. Note the existence of two subpopulations of cells, characterized by different widths, larger or smaller than 1 µm (here separated by the vertical dashed line). Neither the diffusion coefficient along the long cell axis (top, nor along the short cell axis (bottom) vary between these two subpopulations, as shown by the large -values obtained with Kruskal-Wallis H-tests.

(**C**) The values of diffusion coefficient reported in the main text, Fig. 1J, were obtained after applying an “adaptive filter”, as described in the Methods section (y-axes of both plots). These values are compared with either diffusion

coefficients from particles that were not filtered based on their brightness (x-axis in the left plot), or after applying a "fixed filter" with a cutoff value of $\log_{10}$(*brightness*) = 3.5 (x-axis in the right plot).
(**D**) Using the values of *MSD* along the long axis of each cell, $MSD_x$, different diffusion coefficients were estimated: $D_{x,linear}$, from Eq. 1 (y-values on both plots); $D_{x,app}$, from Eq. 2 (x-values, left plot); $\Gamma_x$, from Eq.3 (x-values, right plot).The dashed line represents the diagonal, y = x. Markers show the mean ± standard deviation of n ≥ 3 experimental replicates, except for the "Starved" condition, for which n = 2.
Original units of the diffusion coefficients are $\mu m^2/s$.

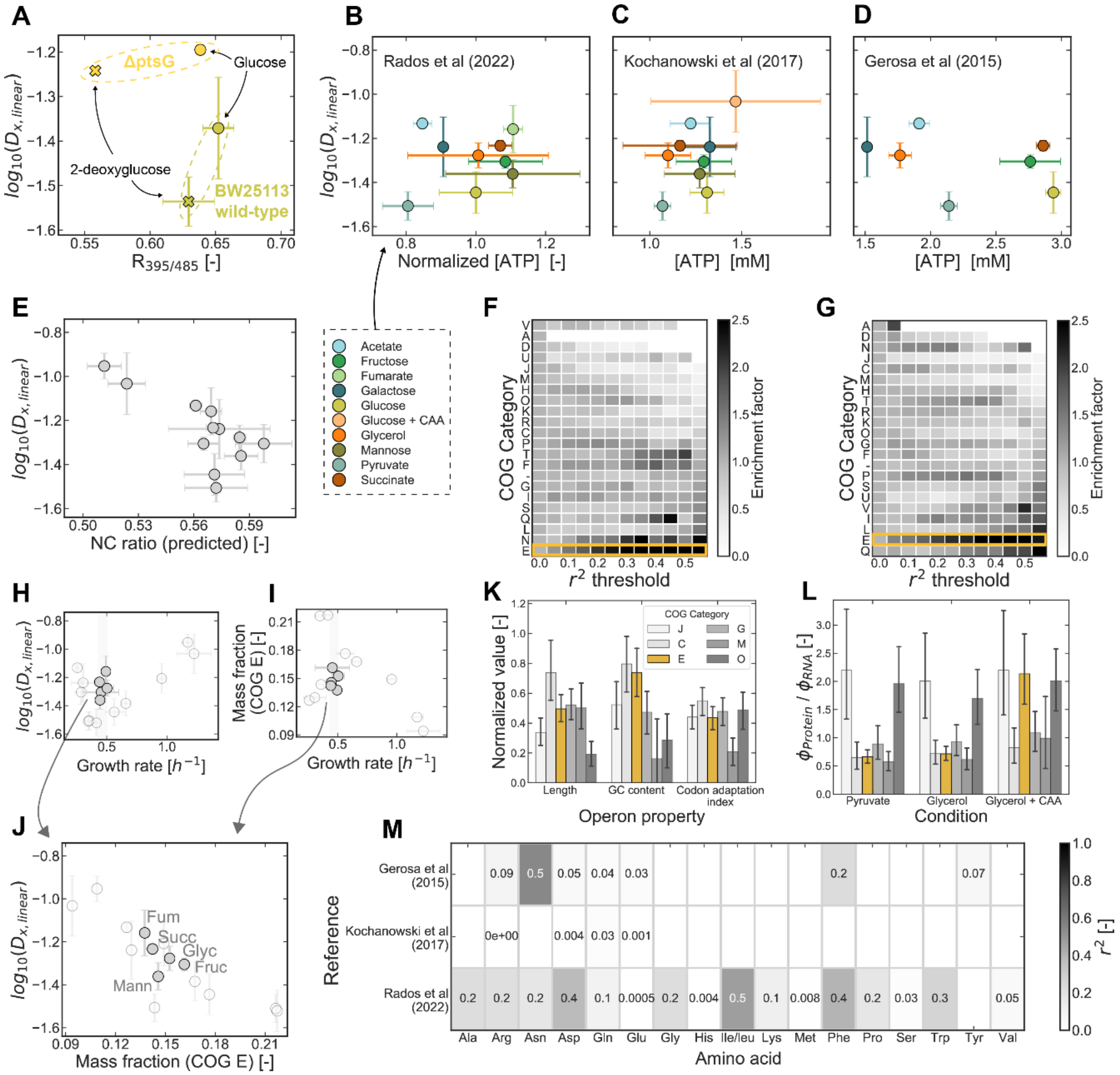


**Figure S2 – Condition-dependence of the diffusion coefficients applies to other metrics of diffusion and is not explained by previously proposed parameters, related to Figure 2.**

(**A**) Diffusion coefficients measured in BW25113 wild-type (dark yellow) and Δ*ptsG* (yellow) strains subjected to different treatments are plotted against a proxy for intracellular pH, namely the ratiometric fluorescence readout of the pHluorin pH-sensor, $R_{395/485}$. Cells were grown in minimal medium with glucose and then washed and resuspended in minimal medium without carbon source. For microscopy imaging, samples were then placed under agarose pads that contained either 5 g/L of glucose (circles), or 5 g/L of 2-deoxyglucose (crosses), a non-metabolizable glucose analogue[128]. This procedure was performed for cells carrying either the pTet-PfVS or pNTR-SD-pHluorin plasmids, for diffusion- and pH-measurements, respectively. Each marker shows the mean ± standard deviation of n = 2 experimental replicates.

(**B**) Diffusion coefficients of the conditions of exponential growth shown in Fig. 1J are plotted against published data on intracellular ATP, from Radoš *et al*[51] (table S2). The data refers to the BW25113 strain, grown in M9 minimal medium. The data obtained are not in absolute concentrations but rather normalized to the concentration measured in glucose-grown cells (dark yellow marker).

(**C**) Same as (**B**), using the data from Kochanowski *et al*[50], taken from its table EV6-1.

(**D**) Same as (**B**), using the data from Gerosa *et al*[49], taken from its data S1. The values, originally reported in units of µmol/gDW, were converted to mM assuming a cell volume of 2.7 µL/mgDW[129]. Error bars for the x-values are the uncertainties reported in the metabolomics data sets. Error bars for the y-values are the standard deviation across experimentally measured values.

(**E**) Diffusion coefficients of the conditions of exponential growth shown in Fig. 1J plotted against the estimated nucleoid-to-cell area ratio, NC. For each growth condition, the average cell area (experimentally measured by microscopy) was used to estimate the nucleoid area. For this estimation, a linear relationship between cell area and nucleoid area was assumed, as previously reported (Fig. 1D in Gray *et al*[30]).

(**F**) Heatmap showing the fractional enrichment of each COG category among the set of proteins with correlation coefficients $r^2 \geq r^2_{threshold}$. These $r^2$ coefficients were obtained by correlating the protein abundance, with the generalized diffusion coefficient, $\Gamma$, obtained from Eq. 3.

(**G**) Similar to (**F**), but with the correlations having been performed with the anomalous diffusion exponent, $\alpha$.

**(H)** Plot of the measured diffusion coefficient of the 40 nm particle against the growth rate, for conditions in which *E. coli* BW25113 displays a narrow range of growth rates, i.e., [0.43, 0.51] $h^{-1}$.

(**I**) Similar to (**H**), showing the mass fraction of COG-E proteins.

(**J**) Plot of the highlighted datapoints in (**H**-**I**). “Fruc”: fructose; “Fum”: fumarate; “Glyc”: glycerol; “Mann”: mannose; “Succ”: succinate.

(**K**) Normalized values of average operon properties do not reveal distinctive characteristics of COG-E transcripts. Each operon was assigned to a COG category based on a “majority” approach. The average length (obtained from Tierrafría *et al*[130]), GC content and codon adaptation index of each COG category were then determined. The latter two properties are proxies for, respectively: the stability of secondary structure formation[131]; and the rate of translation elongation, which describes how long a ribosome remains bound to an mRNA, assuming all mRNAs to have similar length. These values were subjected to a "min-max" normalization, whereby the highest value (across all COG categories) was assigned a value of 1, and the lowest was assigned a value of 0. Note that none of the values displayed is either 0 or 1, indicating that other COG categories (beyond the ones displayed) had the highest/lowest values of the analyzed properties. Only the six COG categories with the highest mass fraction are shown. The bars show the mean ± 95% confidence interval, obtained by bootstrapping.

(**L**) Average ratio between the fractional abundance of a protein, $\phi_{Protein}$ (data from Schmidt *et al*[54]), and the fractional abundance of its corresponding transcript, $\phi_{RNA}$ (data obtained by RNAseq), as a proxy for the number of ribosomes attached to each mRNA molecule. Differently from (**K**), this analysis focuses on the individual protein/transcript level, and not on operons. The values shown are furthermore condition-dependent, varying with the abundances of mRNA and proteins. Only the six COG categories with the highest mass fraction are shown. The bars show the mean ± 95% confidence interval, obtained by bootstrapping. Same colors as in (**K**).

(**M**) Table showing the values of $r^2$ of the correlations between intracellular abundance of various amino acids and the diffusion coefficients measured under the same growth conditions. The metabolomics data was obtained from three publications[49–51]. For some of the amino acids shown, data was available in only one or two of the data sets. Where data was missing, the corresponding rectangle on the table does not display any number. In all cases, the $r^2$ values are below 0.5 (maximum value obtained for asparagine, “Asn”, with Gerosa *et al*’s data).

Original units of the diffusion coefficients are $\mu m^2/s$.

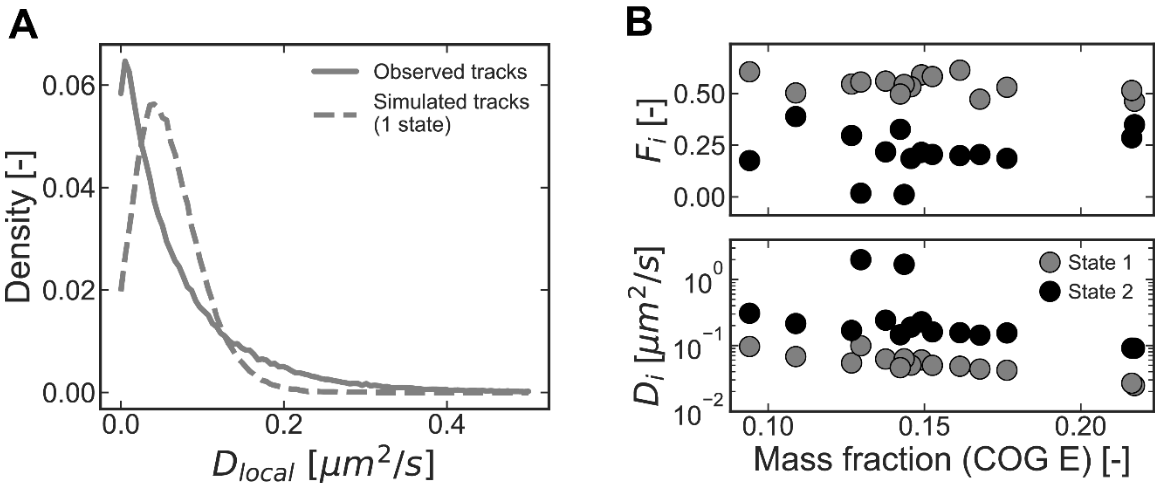


**Figure S3 – Heterogeneity of particle diffusion is captured by multiple diffusive states, related to Figure 3.**

(**A**) Distributions of *local* diffusion coefficients, determined from the *MSDs* in time windows of ten timesteps, using either an experimental data set (growth on medium with galactose; full line), or a simulated set of particle trajectories with a single diffusive state but with the same average diffusion coefficient (dashed line). The simulated trajectories were obtained assuming a 2D-random walk. The larger spread of the former distribution shows that the experimental data cannot be appropriately described by a model with a single diffusive state, and thus a multi-state model better represents the experimental data.

(**B**) Fractions (upper panel) and diffusion coefficients (lower panel) of the "fast" diffusive states, i.e. states 1 and 2, as a function of the mass fraction of COG-E proteins across growth conditions.

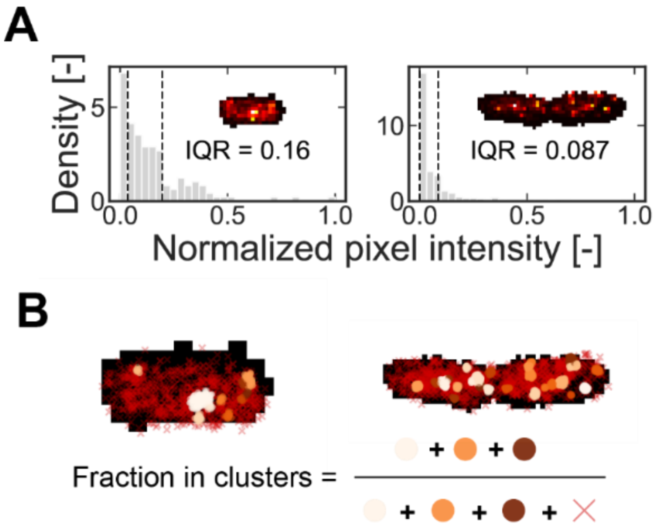


**Figure S4 – Two measures of fluorescence clustering, related to Figure 4.**
Spatial heterogeneity of blinking events of the photoswitchable protein mEos3.2, here shown for two cells expressing ArgD-mEos3.2, was quantified with two metrics:

(**A**) the interquartile range, IQR, of the distribution of normalized pixel intensities in each cell, where the IQR is given by the distance between the two vertical lines. For each pixel, the intensity increases with the number of blinking events detected. Normalizing the values of all pixels inside a cell by the maximum found in the same cell, the IQR is lower for cells that have more heterogeneous distributions of mEos3.2 (e.g., lower IQR for the rightmost cell).

(**B**) the fraction of blinking events assigned to a cluster. Blinking events in each cell were analyzed with a clustering algorithm. Events assigned to a cluster are shown by the colored circles; those not assigned to any cluster are shown with red crosses. The fraction of events assigned to clusters is calculated as shown.

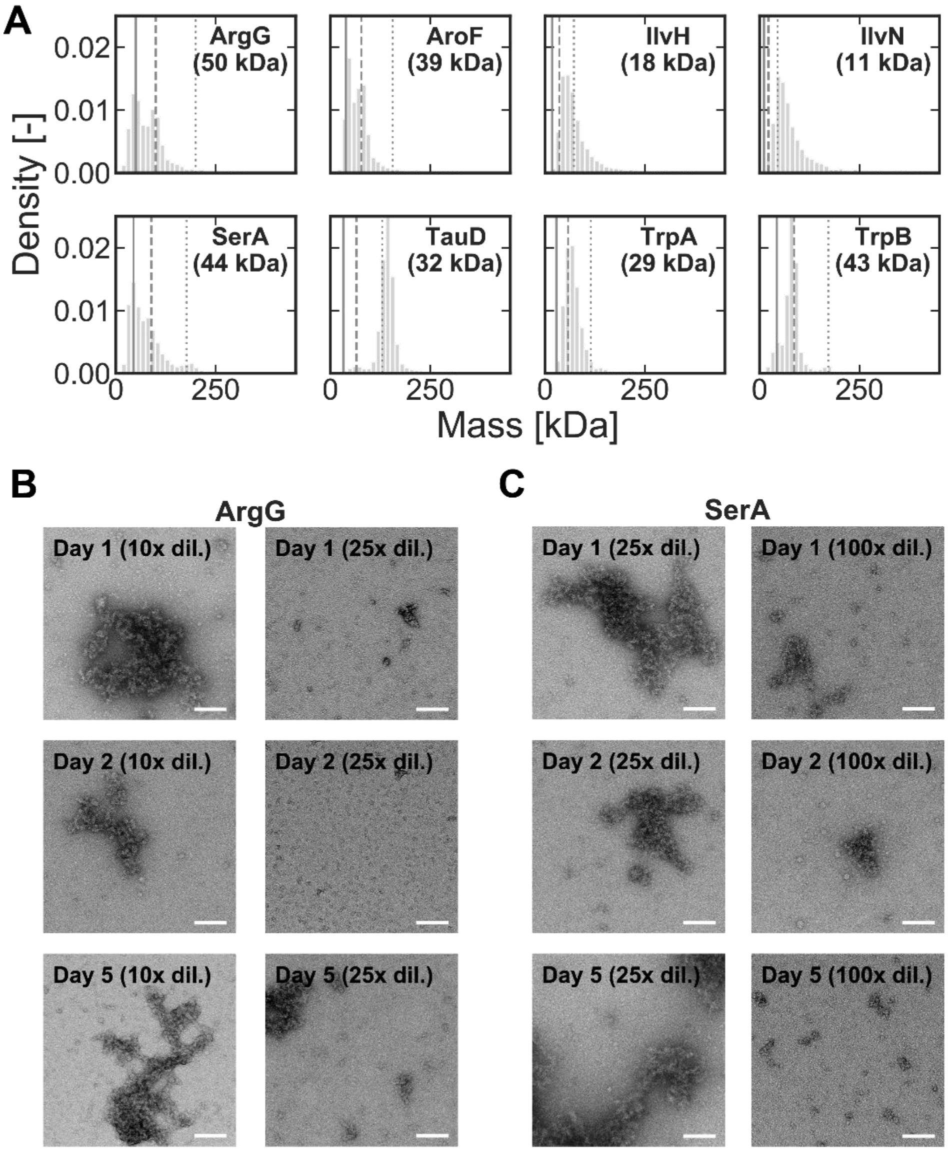


**Figure S5 – In vitro evidence for the formation of agglomerates, related to Figure 5.**

(**A**) Mass distribution, obtained with mass photometry, of eight purified proteins at concentrations of ≤ 50 nM. The eight proteins analyzed are: ArgG, reportedly a homotetramer[132]; AroF, reportedly a homodimer[133]; IlvH, which forms a heterotetramer with IlvI, but is reported as a homodimer in isolation[134]; IlvN, which forms heterotetramer with IlvB, but is reported as a homodimer in isolation[135]; SerA, reportedly a homotetramer[136,137]; TauD, reportedly a homotetramer[138]; TrpA, which forms a heterotetramer with TrpB, although both are reported as homodimers in isolation[139,140]. In each panel, the expected mass of the monomer, homodimer and homotetramer are indicated by the vertical lines (full, dashed and dotted lines, respectively).

(**B**) Negative staining transmission electron microscopy images of ArgG samples at 1-, 2- or 5-days post-purification, during which the protein was kept at 4 °C. For imaging, the stock solution (at 0.6 mg/mL) was diluted with the dilution factors indicated on each panel (10x or 25x). Scale bar: 100 nm. Sample “Day 1 (10x dil.)” is also shown in Fig. 5I.

(**C**) Same as (**B**) for the protein SerA, from a stock solution at 1.5 mg/mL, and pre-diluted by a factor of 25x or 100x.

### Supplemental information

**Table S1 – Features used to train the first random forest classifier (Model #1), related to Figure 5.** Features describing each protein were obtained from different sources [S1, S2] or determined with computational tools that rely on either sequence or structure data (*BioPython, IUPred2A, MDTraj, MolPatch, NetSurfP3.0, PyBioMed*). From an initial set of 138 features, we removed: (i) one per pair of features with a Pearson correlation coefficient higher than 0.8, and (ii) features with a Variance Inflation Factor higher than 5. Here, the set of remaining 76 non-collinear features which used with Model #1 (Fig. 5B) is shown.

| Based on | Feature name | Explanation |
|---|---|---|
| - | avg_abundance | Average protein abundance reported by Schmidt *et al* (2016) across the growth conditions used in this study. |
| | noInteractions | Number of protein-protein interactions reported on BioGRID. Obtained from *UniProt*. |
| | noPTMs | Number of post-translational modifications. Obtained from *UniProt*. |
| | cytoplasmic_True | Binary feature indicating if a protein's location is described to be the cytoplasm. Obtained from *UniProt*. |
| | dnaBinding_True | Binary features indicating if a is known to bind to DNA, RNA, metal ions or ATP. Obtained from *UniProt*. |
| | rnaBinding_True | |
| | ionBinding_True | |
| | atpBinding_True | |
| Sequence | _biopython_pi | Estimated isoelectric point. Obtained with *Biopython*. |
| | _biopython_aacomp_D | Fraction of each amino acid in the protein sequence. Obtained with *Biopython*. |
| | _biopython_aacomp_H | |
| | _biopython_aacomp_I | |
| | _biopython_aacomp_L | |
| | _biopython_aacomp_M | |
| | _biopython_aacomp_N | |
| | _biopython_aacomp_P | |
| | _biopython_aacomp_Q | |
| | _biopython_aacomp_R | |
| | _biopython_aacomp_S | |
| | _biopython_aacomp_T | |
| | _biopython_aacomp_V | |
| | _biopython_aacomp_W | |
| | _biopython_aacomp_Y | |
| | _pybiomed_ctd__SolventAccessibility**T**13 | Transition and distribution descriptors ("T" and "D", respectively; bold underlined) of a protein sequence in terms of the solvent accessibilities of its residues. The following encoding of residues applies: buried ("1"), exposed ("2") and intermediate exposure ("3"). "T" counts the occurrence where residues of one group are immediately followed by residues of another group (e.g. "T13" = transition between residues of types "1" and "3"). |
| | _pybiomed_ctd__SolventAccessibility**D**1025 | |
| | _pybiomed_ctd__SolventAccessibility**D**3025 | |
| | _pybiomed_ctd__SolventAccessibility**D**3075 | |

| | | |
|---|---|---|
| | | "D" describes the fraction of residues of one group in the first 25 or 75% of the sequence (e.g. "D1025" = fraction of residues "1" among the first 25% of the entire sequence).<br>Obtained with *PyBioMed*. |
| | _pybiomed_ctd__SecondaryStr**T**12 | Same as above, describing protein sequence in terms of the secondary structures in which the aminoacids are typically found. The following encoding applies: helix ("1"), strand ("2") and coil ("3"). |
| | _pybiomed_ctd__SecondaryStr**T**13 | |
| | _pybiomed_ctd__SecondaryStr**T**23 | |
| | _pybiomed_ctd__SecondaryStr**D**1025 | |
| | _pybiomed_ctd__SecondaryStr**D**1075 | |
| | _pybiomed_ctd__SecondaryStr**D**2025 | |
| | _pybiomed_ctd__SecondaryStr**D**2075 | |
| | _pybiomed_ctd__SecondaryStr**D**3025 | |
| | _pybiomed_ctd__SecondaryStr**D**3075 | |
| | _pybiomed_ctd__Charge**T**13 | Same as above, describing protein sequence in terms of the charge of the residues. The following encoding applies: positive ("1"), neutral ("2") and negative ("3"). |
| | _pybiomed_ctd__Charge**D**1025 | |
| | _pybiomed_ctd__Charge**D**1075 | |
| | _pybiomed_ctd__Charge**D**2025 | |
| | _pybiomed_ctd__Charge**D**2075 | |
| | _pybiomed_ctd__Charge**D**3025 | |
| | _pybiomed_ctd__Charge**D**3075 | |
| | _pybiomed_ctd__Polarity**T**12 | Same as above, describing protein sequence in terms of residue polarity. The following encoding applies: low polarity, 4.9 – 6.2 ("1"), medium polarity, 8.0-9.2 ("2"), and high polarity, 10.4 – 13.0 ("3"). |
| | _pybiomed_ctd__Polarity**T**13 | |
| | _pybiomed_ctd__Polarity**T**23 | |
| | _pybiomed_ctd__Polarity**D**2025 | |
| | _pybiomed_ctd__Polarity**D**2075 | |
| | _pybiomed_ctd__Polarity**D**3025 | |
| | _pybiomed_ctd__Polarity**D**3075 | |
| | _pybiomed_ctd__NormalizedVDWV**T**12 | Same as above, describing protein sequence in terms of the normalized van der Waals volumes of the residues. The following encoding applies: low, 0 – 2.78 ("1"), medium, 2.95 – 4.0 ("2"), and high, 4.03 – 8.08 ("3"). |
| | _pybiomed_ctd__NormalizedVDWV**T**13 | |
| | _pybiomed_ctd__NormalizedVDWV**T**23 | |
| | _pybiomed_ctd__NormalizedVDWV**D**1025 | |
| | _pybiomed_ctd__NormalizedVDWV**D**1075 | |
| | _pybiomed_ctd__NormalizedVDWV**D**2025 | |
| | _pybiomed_ctd__NormalizedVDWV**D**2075 | |
| | _pybiomed_ctd__NormalizedVDWV**D**3025 | |
| | _pybiomed_ctd__NormalizedVDWV**D**3075 | |
| | _pybiomed_ctd__Hydrophobicity**T**12 | Same as above, describing protein sequence in terms of the aminoacids hydrophobicity. The following encoding applies: polar ("1"), neutral ("2") and hydrophobic ("3"). |
| | _pybiomed_ctd__Hydrophobicity**T**23 | |
| | _pybiomed_ctd__Hydrophobicity**D**1025 | |
| | _pybiomed_ctd__Hydrophobicity**D**1075 | |
| | _pybiomed_ctd__Hydrophobicity**D**3025 | |
| | _pybiomed_ctd__Hydrophobicity**D**3075 | |

| | | |
|---|---|---|
| | _iupred_max_IUPredscore | Highest value, across all amino acids in the sequence, of the disorder score predicted with *IUPred2A*. |
| | _iupred_ noDomains | Number of “disordered domains”, which are defined as stretches of at least 10 consecutive amino acids with a disorder score higher than 0.5, as predicted with *IUPred2A*. |
| | _iupred_max_DomainLength | Number of amino acids in the longest “disordered domain”, defined as above. Determined using the ouput of *IUPred2A*. |
| | _nsp3_disorder | Average, over all amino acids in the protein sequence, of the disorder score predicted with *NetSurfP3.0*. |
| | _nsp3_asa | Sum, over all amino acids in the protein sequence, of the solvent accessible area predicted with *NetSurfP3.0*. |
| | _nsp3_H | Fraction of amino acids that are predicted to form alpha-helices, according *NetSurfP3.0*. |
| 3D structure | _mdtraj_acilindricity | Acilindricity of the *AlphaFold*-predicted protein structure. Determined with *MDTraj*. |
| | _mdtraj_asphericity | Asphericity of the *AlphaFold*-predicted protein structure. Determined with *MDTraj*. |
| | _mdtraj_anisotropy | Shape anisotropy of the *AlphaFold*-predicted protein structure. Determined with *MDTraj*. |
| 3D structure + Sequence | hydropathy_surface_density | Average hydropathy score (Kyte-Doolittle scale) across amino acids, weighted by their relative solvent accessible area. |
| | _molpatch_patchSize | Area of the largest hydrophobic patch on the surface of the *AlphaFold*-predicted protein structure. Determined with *MolPatch*. |

**Table S2 – Selected features used to train the second random forest classifier (Model #2), related to Figure 5.** Set of 17 features, mostly describing hydrophobicity and three-dimensional protein shape. The six features marked with an asterisk (*) were excluded during the automated removal of collinear features. Model #2 (Fig. 5D) was thus trained with the remaining 11 features.

| Based on | Feature name | Explanation |
|---|---|---|
| - | avg_abundance | Average protein abundance reported by Schmidt *et al* (2016) across the growth conditions used in this study. |
| | cytoplasmic_True | Binary feature indicating if a protein's location is reported to be the cytoplasm. Obtained from *UniProt*. |
| Sequence | Length (*) | Number of amino acids in a protein sequence. |
| | hydropathy_mean_overall | Average, over all amino acids in a sequence, of the hydropathy score (Kyte-Doolittle scale). |
| 3D structure + Sequence | hydropathy_mean_exposedOnly | Average, over all amino acids with a relative solvent-accessible area >25%, of the hydropathy score (Kyte-Doolittle scale). The relative solvent accessible area of each amino acid was determined with *MDTraj* using the *AlphaFold*-predicted protein structures. |
| | SASA/length | Total solvent accessible surface area, $SASA_{total}$, normalized by the sequence length. Determined with *MDTraj*. |
| | surface_area_hydrophobic (*) | Total solvent accessible area of all hydrophobic amino acids (A,C,F,I,L,M,V,W,Y), $SASA_{hydrophobic}$. Determined with *MDTraj*. |
| | rel_surface_area_hydrophobic (*) | Ratio between $SASA_{hydrophobic}$ and $SASA_{total.}$ |
| 3D structure | _mdtraj_rg (*) | Shape descriptors of the *AlphaFold*-predicted protein structures: radius of gyration, asphericity and relative shape anisotropy. Determined with *MDTraj*. |
| | _mdtraj_asphericity | |
| | _mdtraj_anisotropy | |
| | _molpatch_patchSize_0 | Areas of the three largest patches of hydrophobic amino acids on the surface on the surface of the *AlphaFold*-predicted protein structure. Determined with *MolPatch*. |
| | _molpatch_patchSize_1 (*) | |
| | _molpatch_patchSize_2 | |
| | _molpatch_patchSize_0/SASA (*) | Same as above, normalized by $SASA_{total}$. |
| | _molpatch_patchSize_1/SASA | |
| | _molpatch_patchSize_2/SASA | |

**Table S3 – Concentrations of selected proteins in *E. coli* and in vitro experiments described in this work, related to Figures 5 and S5.**

The in vivo abundances were obtained from Schmidt et al [S1], Table S6, after averaging across the ten conditions of exponential growth shown in Fig. 1B. The protein concentrations used for mass photometry and transmission electron microscopy, TEM, were determined by absorbance at 280 nm. Note that the reported protein abundances, *p* (in copies/cell), are approximately similar to the volumetric concentrations, *c* (in nM), given that *c = p/V*, where *V* is the cell volume (approximately equal to 1 μm$^3$/cell [S3]), and $c = \left(\frac{p}{V}\frac{copies}{\mu m^3}\right) * \left(\frac{1}{10^{-15}}\frac{\mu m^3}{L}\right) * \left(\frac{1\,nmol}{6.022\times10^{23}\,copies}\right) = \left(\frac{p}{V}\times 1.66\right) nM$.

| Protein | In vivo abundance [copies/cell ≈ nM] | Concentration used for mass photometry [nM] | Concentration used for negative staining TEM [nM] |
|---|---|---|---|
| ArgD | 2 242 | ~30 | --- |
| ArgE | 1 239 | ~50 | ~1 071 |
| ArgG | 3 942 | ~10 | ~1 202 |
| AroF | 1 799 | ~50 | ~737 |
| IlvI | 352 | ~20 | ~645 |
| TrpB | 3 351 | ~20 | ~465 |
| TyrA | 1 045 | ~50 | --- |
| SerA | 5 425 | ~10 | ~1 364 |
| Notes | | Refers to the data shown in (Fig. 5G and S5A) | Refers to the images shown in (Fig. 5I) |

#### *Supplemental references*

[S1] Schmidt, A., Kochanowski, K., Vedelaar, S., Ahrné, E., Volkmer, B., Callipo, L., Knoops, K., Bauer, M., Aebersold, R., and Heinemann, M. (2016). The quantitative and condition-dependent Escherichia coli proteome. Nat. Biotechnol. *34*, 104–110. https://doi.org/10.1038/nbt.3418.

[S2] The Uniprot Consortium (2023). UniProt: the Universal Protein Knowledgebase in 2023. Nucleic Acids Res. *51*, D523–D531.

[S3] Milo, R., and Phillips, R. (2015). Cell Biology by the Numbers 1st ed. (Garland Science) https://doi.org/https://doi.org/10.1201/9780429258770.
